# Structural Basis for Childhood Antibody Recognition of The Human Metapneumovirus Fusion Protein

**DOI:** 10.1101/2025.08.29.673069

**Authors:** Ahmed Magdy Khalil, Behrouz Ghazi Esfahani, Rose J. Miller, Jiachen Huang, Guillermina Kuan, Angel Balmaseda, Aubree Gordon, Jarrod J. Mousa

## Abstract

Human metapneumovirus (hMPV) is a significant cause of acute respiratory illness in children and adults, with the majority of children being seropositive for hMPV by five years of age. Infants, older adults, and immunocompromised individuals are more susceptible to severe hMPV infections that can lead to hospitalization and death. The hMPV fusion (F) protein is the sole target of neutralizing antibodies, and while the most common neutralizing epitopes on the hMPV F protein targeted by B cells in hMPV-infected adults have been previously determined, the antibody response in hMPV-infected children remains undefined. We isolated a panel of human monoclonal antibodies (mAbs) from children previously infected with hMPV (MPV498, MPV499, MPV510, MPV511, and MPV513), and the mAbs were assessed for binding avidity, neutralization potency, epitope specificity, and *in vivo* efficacy. All mAbs were neutralizing, and epitope binning revealed the presence of four different epitopes targeted by the mAbs. We determined the cryo-EM structures of four mAbs in complex with the hMPV F protein, which revealed epitopes located on the hMPV F trimer surface as well as an intratrimer epitope located completely within the hMPV F trimer interface. Furthermore, we determined the prophylactic efficacy of the mAbs in protection against hMPV challenge in mice. Overall, our data reveal new insights into the immunodominant antigenic epitopes on the hMPV F protein in children and identify new mAb therapies for hMPV F disease prevention.

## Introduction

Human metapneumovirus (hMPV) is an enveloped, non-segmented, single-stranded, negative-sense RNA virus that belongs to the family Pneumoviridae and is closely related to respiratory syncytial virus (RSV)^1^. Indeed, hMPV is a leading cause of acute respiratory infection (ARI) in children and older adults^2, 3^. Although the virus was first identified in 2001 in the Netherlands from children with respiratory tract infections, serological analysis has clarified that hMPV has been circulating in human populations for at least seven decades^4^. By the age of five years, nearly all children are seropositive for hMPV, which implies the wide prevalence of hMPV infections in early life^5^. The clinical features of hMPV infection encompass ARIs and non-classical ARIs such as aggravation of chronic diseases, particularly in elderly and immunocompromised patients (e.g., asthma or chronic obstructive pulmonary disease)^6, 7^. In the United States, the burden of hMPV infection in children under the age of five years includes an annual hospitalization of ∼20,000 children, with more than 260,000 emergency visits and almost ∼1 million outpatient visits^5^, and ∼122,000 annual hospitalizations in adults aged 65 and older with a mortality rate as high as 4.8% in one reported study^8, 9^. Also, recent reports of an increase in the number of hMPV infections in northern Chinese provinces was reported in late 2024, which heightened public health awareness of hMPV^10, 11^. However, there are currently no FDA-approved vaccines or antiviral therapies available to protect against or to treat hMPV infection.

There are three proteins on the envelope of hMPV: the small hydrophobic (SH) protein, the attachment (G) protein, and the fusion (F) protein. Among these proteins, only the F protein elicits neutralizing antibodies, while immunization with soluble hMPV G protein was shown to be immunogenic but nonprotective in cotton rats^12^, standing in contrast to RSV, where both RSV G and RSV F proteins elicit neutralizing antibodies and protect against viral replication *in vivo*^13, 14^. The hMPV F protein is a class I trimeric membrane-bound glycoprotein that mediates the fusion of the viral envelope to the host cell membrane during virus infection. Heparan sulfate and RGD-binding integrins have been previously discovered as host cell receptors for the hMPV F protein^15–17^. Initially, the F protein is synthesized as a fusogenically inactive precursor (F0) that is subsequently cleaved by host cell serine proteases, e.g., TMPRSS2, into disulfide-linked F_2_ and F_1_ subunits that undergo conformational changes to facilitate membrane fusion and subsequent virus uncoating to release viral RNA genome into the cytoplasm of the host cell^18, 19^.

Advancements in structure-based protein engineering have led to the determination of X-ray crystal structures of the hMPV F protein in both pre-fusion and post-fusion conformations, and these protein constructs have been leveraged to determine the major immunogenic epitopes of neutralizing antibodies in adults who are seropositive for hMPV^17, 20–23^. Unlike RSV, where the majority of antibodies target the pre-fusion conformation of the RSV F protein^24–26^, the majority of hMPV F antibodies discovered from humans bind both the pre-fusion and post-fusion conformations^17, 22, 27^. Monoclonal antibodies (mAbs) identified were mapped to antigenic site Ø^27, 28^, the DS7 antigenic site (site I)^29^, antigenic site III^30–32^, antigenic site IV^23, 33^, and antigenic site V^34^. Additionally, two epitopes located within the trimeric interface of the hMPV F protein have been identified^35, 36^.

Similar to RSV, formalin-inactivated hMPV vaccines result in vaccine-enhanced disease upon subsequent exposure to virus infection in animal models^37–40^. Formalin-inactivation of RSV has been demonstrated to cause changes in the F protein from the pre-fusion to the post-fusion conformation^38^. As pre-fusion stabilization of the RSV F protein has led to the success of vaccine approval and disease reduction in humans^41, 42^, presumably this success would translate to RSV-naïve children. However, signals of vaccine-enhanced disease were recently observed in a clinical trial with the mRNA-LNP-based vaccine against RSV/hMPV (mRNA-1345 and mRNA-1365, NCT05743881) in infants and children younger than two years old. Meanwhile, additional candidates including an mRNA-LNP-based vaccine (NCT06686654), a protein subunit-based vaccine (NCT06984094), and virus-like particle protein vaccine (AstraZeneca/Icosavax) are undergoing clinical trials for efficacy against hMPV.

Human mAbs continue to be a growing class of drugs, in part due to their high degree of specificity, limited off-target effects, and superb safety profile^43^. For instance, nirsevimab and clesrovimab, FDA-approved anti-RSV mAbs, have recently been approved to protect infants from RSV infection^44^. Additionally, palivizumab, a humanized mAb, has been used for several years to protect high-risk infant groups from RSV infection^45^. While the B cell response to RSV infection in adults^25, 31, 46–49^ and children^50^ has been well studied, only studies on hMPV infection in adults^27, 47, 48, 51–54^ have been conducted, and with the recent results from the Moderna mRNA-LNP vaccine trial, the lack of knowledge of the immunological profile of hMPV in children remains a barrier to the development of protective hMPV vaccines^55^.

Herein, we isolated five mAbs (MPV498, MPV499, MPV510, MPV511, and MPV513) from children infected with hMPV. The five mAbs showed strong binding avidity and varied neutralizing activity against hMPV strains of different genotypes. Epitope binning revealed four epitopes targeted by the five mAbs. We determined the cryo-EM structure of four mAbs in complex with the hMPV F protein, which revealed epitopes located on the hMPV F trimer surface as well as intratrimer epitopes. Furthermore, we revealed the prophylactic efficacy of the five mAbs in protection against hMPV infection in mice.

## Results

### Isolation of hMPV-specific mAbs

We isolated five mAbs from hMPV-infected children who were diagnosed during a cohort study from 2011 to 2016 in Managua, Nicaragua^56^ **(Table S1)** using hMPV B2 F protein^17^ as the antigen bait. mAb MPV498 was generated using human hybridoma technology^30^, while the remaining four mAbs (MPV499, MPV510, MPV511, and MPV513) were derived from antigen-specific single B cell sorting^57, 58^. The antibody-encoding genes were sequenced **(Table S2)**, and analysis with IMGT^59^ indicated the usage of the variable heavy (VH) 4 gene family for mAbs MPV498, MPV499, MPV510, and MPV511, whereas mAb MPV513 belonged to the VH5 gene family. Also, mAbs MPV499, MPV510, and MPV513 shared the same diversity heavy (DH) 3 and joining heavy (JH) 4 gene families, while mAbs MPV498 and MPV511 utilized DH5-JH5 and DH2–JH6 gene families, respectively **(Table 1)**. Furthermore, all five mAbs utilize the kappa light chain isotype (VK) 1 gene family: mAbs MPV498, MPV499, MPV10, and MPV513 utilize the VK1-39 gene, while mAb MPV511 utilizes the VK1-5 gene. Also, mAbs MPV498 and MPV513 utilize the same JL2 gene, mAbs MPV510 and MPV511 utilize the JL1 gene, and mAb MPV499 belongs to the JL4 gene family **(Table 1)**. The lengths of the heavy chain (HC) junction regions were 15 amino acids for mAbs MPV499, MPV510, and MPV513, 17 amino acids for mAb MPV511, and 20 amino acids for mAb MPV498. Whereas the lengths of junctions in the light chain (LC) were 11 amino acids for all mAbs except MPV510 mAb, which was 12 amino acids long **(Table S2)**. The percent identities of the variable genes to the germline sequence ranged from 87.72−98.25% (94.5% average) for the HC and 93.91−97.85% (95.79% average) for the LC, which are comparable to the variable gene identities of previously reported hMPV F-specific mAbs^51^ **(Table 1)**.

**Table 1.** Gene usage and percent identity of the hMPV F-specific mAbs^a^.

| Table 1. Gene usage and percent identity of the hMPV F-specific mAbs<sup>601</sup> |  |  |  |  |  |
| --- | --- | --- | --- | --- | --- |
| mAb | HC |  |  | LC |  |
|  | VH (% identity) | DH | JH | VL (% identity) | JL<sup>602</sup> |
| **MPV498** | VH4-4 (98.25) | D5-12 | J5 | VK1-39 (97.38) | J2<sup>603</sup> |
| **MPV499** | VH4-31 (96.91) | D3-3 | J4 | VK1-39 (94.12) | J4 |
| **MPV510** | VH4-30 (92.78) | D3-10 | J4 | VK1-39 (95.7) | J1 |
| **MPV511** | VH4-34 (87.72) | D2-21 | J6 | VK1-5 (93.91) | J1 |
| **MPV513** | VH5-10 (96.88) | D3-3 | J4 | VK1-39 (97.85) | J2 |
| <sup>a</sup>Analysis was performed using IMGT/V-Quest. |  |  |  |  |  |

### Binding and neutralizing properties of the isolated mAbs

The binding properties of the five mAbs were assessed by enzyme-linked immunosorbent assay (ELISA) using a panel of hMPV F proteins comprising the four subgroups of hMPV (A1, A2, B1, and B2), pre-fusion stabilized hMPV F DsCavEs2^60^ protein, and the post-fusion hMPV B2 F protein as previously described^17^. All five mAbs bound to the four subgroups of the hMPV F protein as well as the pre-fusion stabilized hMPV F DsCavEs2 protein with an EC_50_<1.0 μg/mL, whereas only mAbs MPV498 (EC_50_ = 0.16 μg/mL) and MPV511 (EC_50_ = 0.05 μg/mL) showed binding affinity to the post-fusion hMPV B2 F protein **(Figure 1A)**, which indicates the pre-fusion binding preference of the isolated mAbs. The neutralizing activity of the five mAbs was determined by plaque-reduction neutralization assay using representative viruses from hMPV genotype A (strain CAN/97-83, subgroup A2) and genotype B (strain TN/93-32, subgroup B2). All five mAbs had neutralizing activity against both hMPV genotypes, with MPV511 being the most potent mAb with an IC_50_ of 0.35 and 0.02 μg/mL against both hMPV CAN/97-83 and TN/93-32 viruses, respectively. Notably, MPV499, MPV510, and MPV511 mAbs showed more potent neutralizing activity against TN/93-32 virus, mAb MPV498 showed more potent neutralizing activity against CAN/97-83, and mAb MPV513 had similar neutralizing activity against both viruses **(Figure 1B)**.

**Figure 1.**
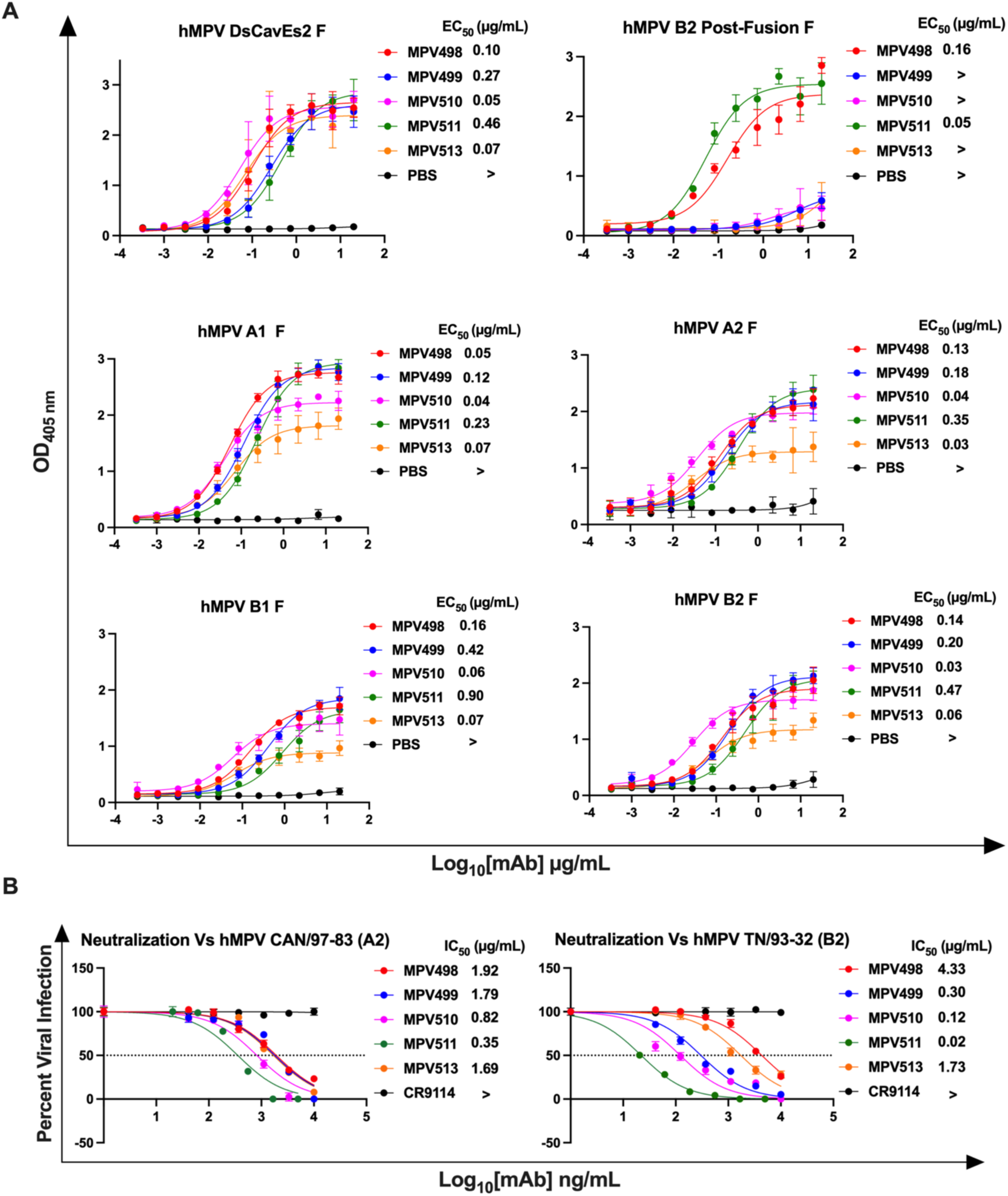
ELISA binding and virus neutralization of the five isolated mAbs. (A) ELISA binding curves of isolated mAbs against different hMPV proteins. EC_50_ values correspond to the concentration at which the half-maximum signal was obtained in ELISA based on the optical density at 405 nm (> indicates signal at highest concentration did not reach 1.0). Each point represents the average of data from four technical replicates, and error bars represent the standard deviation. Data are representative of two biological replicates. (B) Plaque reduction neutralization assays of the five isolated mAbs against two different strains of hMPV. The IC_50_ corresponds to the mAb concentration at which 50% plaque reduction was observed (> indicates no reduction > 50% at the highest concentration). Each point represents the average of data from three technical replicates, and error bars represent the standard deviation. Data are representative of two biological replicates.

### Epitope mapping of the isolated mAbs

To determine the hMPV F binding epitopes for the five isolated mAbs, an epitope binning experiment was conducted in a pairwise combinatorial manner, where each mAb was tested against itself and other mAbs using biolayer interferometry (BLI) as previously described **(Figure 2)**^30, 35, 61, 62^. Control mAbs targeting the hMPV F protein, including MPE8^31^ and MPV364^54^ (site III), MPV481^51^ (site IV), MPV467^51^ (site V), MPV458^35^ (trimeric interface amino acids 66-87 of the F2 region), and MPV196^54^ (DS7 antigenic site I) were included to determine the competition profiles for the tested mAbs. Briefly, anti-histidine tag biosensors were loaded with the pre-fusion stabilized hMPV DsCavEs2 F protein, associated first with either control mAbs or test mAbs, and then exposed to the secondary mAb **(Figure 2A)**. In the first experiment, where control mAbs were loaded first and then exposed to the tested mAbs, the primary binding of mAb MPV458 to the 66-87 trimeric interface epitope blocked the binding of mAb MPV498 to its epitope. mAb MPV511 mapped to an overlapping epitope between antigenic sites III and DS7 (site I), where it competed with mAbs MPE8 and MPV196, respectively, which is similar to our previously identified binding approach for mAb MPV196, as antigenic site III lies in close proximity to site I^54^. mAbs MPV499, MPV510, and MPV513 did not compete with any of the control mAbs and only showed partial competition with antigenic site III-targeted mAbs MPE8 and MPV364, suggesting their potential binding to newly unidentified epitopes. Additionally, mAbs MPV499 and MPV513 showed the same binding profiles with partial competition to the 66-87 trimeric interface and DS7 antigenic sites, indicating that these mAbs might share overlapping epitopes **(Figure 2B)**. In the second experiment where tested mAbs were loaded first and then followed by the control mAbs, mAb MPV498 partially competed with mAb MPV458 on the 66-87 epitope, indicating that binding epitopes of both mAbs might overlap, where the binding of mAb MPV498 does not completely block the access of mAb MPV458. The primary binding of either MPV499, MPV510, or MPV513 mAbs completely blocked the binding of the trimeric interface mAb MPV458, which implies possible sharing of a similar epitope, or binding of any of these mAbs disrupts the binding epitope of mAb MPV458. Also, four of the tested mAbs (MPV498, MPV499, MPV510, and MPV513) were grouped in one group where the binding of any of the four mAbs completely or partially blocked the binding of the three others, unlike the binding profile of mAb MPV511, which was unique **(Figure 2B)**.

**Figure 2.**
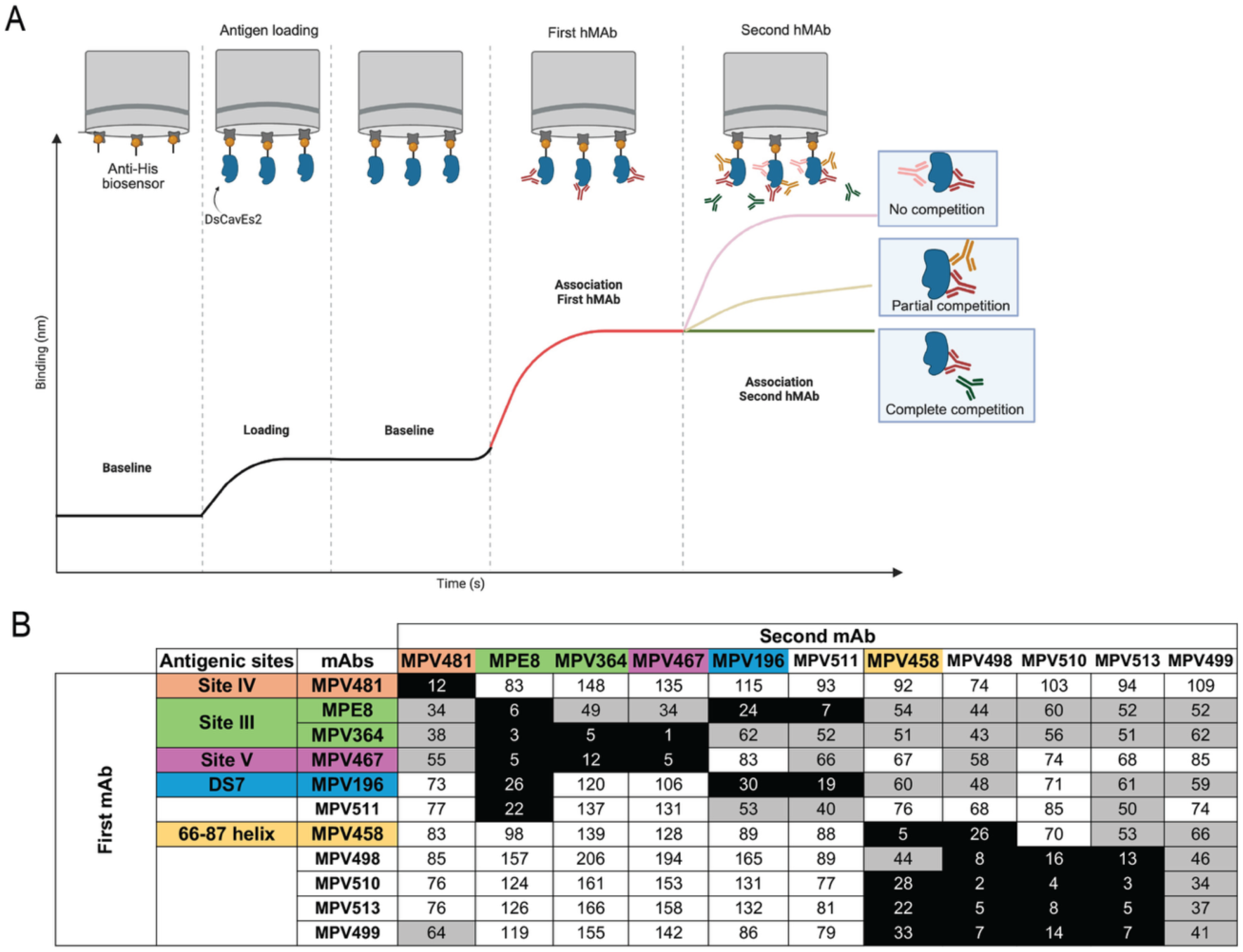
Biolayer Interferometry (BLI) epitope binning of the isolated mAbs against the hMPV DsCav-Es2 F protein. (A) Diagram demonstrating the epitope binning protocol using competitive BLI, where the antigen is loaded first before sequential exposures to the mAbs to determine whether the first mAb blocks binding of the second mAb. (B) A heat map showing the results from a binning experiment with the five mAbs (MPV498, MPV499, MPV510, and MPV513) against control mAbs MPV458, MPV481, MPE8, MPV364, MPV467, and MPV196. (Black [complete competition, ≤33%], gray [partial competition, 34-66%], and white [no competition, ≥67%]). The formula to calculate the competition percentage is %Competition = (Signal from first mAb with second mAb / Signal from first mAb) X 100%.

### Cryo-EM structures of the isolated mAbs

In order to identify the binding sites of the new mAbs, we utilized single particle cryo-electron microscopy. We determined the structures of fragment antigen-binding (Fab) regions from four mAbs (MPV498, MPV499, MPV510, and MPV513) in complex with different conformations of the hMPV F protein. We determined the structure of Fab MPV498 in complex with the hMPV B2 post-fusion F protein using cryo-EM to a global resolution of 2.61 Å obtained from 111,000 particles **(Figure 3, Figure S1, Figure S2A, Table S3)**. The cryo-EM structure reveals that MPV498 Fab makes numerous interactions with multiple residues on the surface of the hMPV B2 post-fusion F protein **(Figure 3A and Figure 3B)**. SER36 at the complementarity-determining region 1 (CDR1) of the MPV498 HC interacts with THR49 (located at the N-terminal of F2 subunit) of the hMPV F protein via a hydrogen bond **(Figure 3B)**, whereas other HC amino acid residues formed several hydrogen bonds with multiple residues located near to or at the Heptad Repeat B (HRB) region at the C-terminal end of the F1 subunit: THR111 (CDR3) with TYR233 (located central at the HRA-HRB loop), SER59 and ASN64 (CDR2) with GLU431 (HRB), TYR38 (CDR1) with GLY432 (HRB), and both SER107 and ARG108 (CDR3) with GLN434 (HRB) **(Figure 3B)**. The MPV498 LC forms two hydrogen bonds through amino acids TYR38 (CDR1) and TYR108 (CDR3) with ARG396 located in the C-terminal end of the F1 subunit (near HRB), likely anchoring the antibody to the base of the hMPV F trimer **(Figure 3B)**. The LC of MPV498 Fab touches only ARG396 within the epitope of antigenic site IV (**Figure 3B).** Overall, the MPV498 interactive amino acid residues with the post-fusion hMPV F protein constitute a new unidentified antigenic epitope (referred to here as hMPV antigenic site VI) that is not reported at any of the previously known hMPV antigenic sites^63^, and is unique from other epitopes near the area such as site II (residues 254-277 located at HRA)^64^ and site IV (residues 391-409, hMPV F numbering located just before HRB) targeted by RSV and hMPV cross-reactive mAbs^23, 47^. Although we did not have the cryo-EM structure for the MPV498 with hMPV F in the pre-fusion conformation, we confirmed the binding ability of mAb MPV498 to the pre-fusion F as well **(Figure 1A**, **Figure 2B, and Figure S3).** mAb MPV498 showed binding to the hMPV DsCavEs2 F protein in ELISA **(Figure 1A)** and using both competitive epitope-binning **(Figure 2B)** and kinetics **(Figure S3)** BLI. Notably, most of the MPV498-binding epitope in the post-fusion conformation (GLU431, GLY432, and GLN434) is buried in the pre-fusion state of the DsCavEs2 F protein due to the conformational changes **(Figure 3C)**, indicating that its binding epitope to the pre-fusion F might be variable.

**Figure 3.**
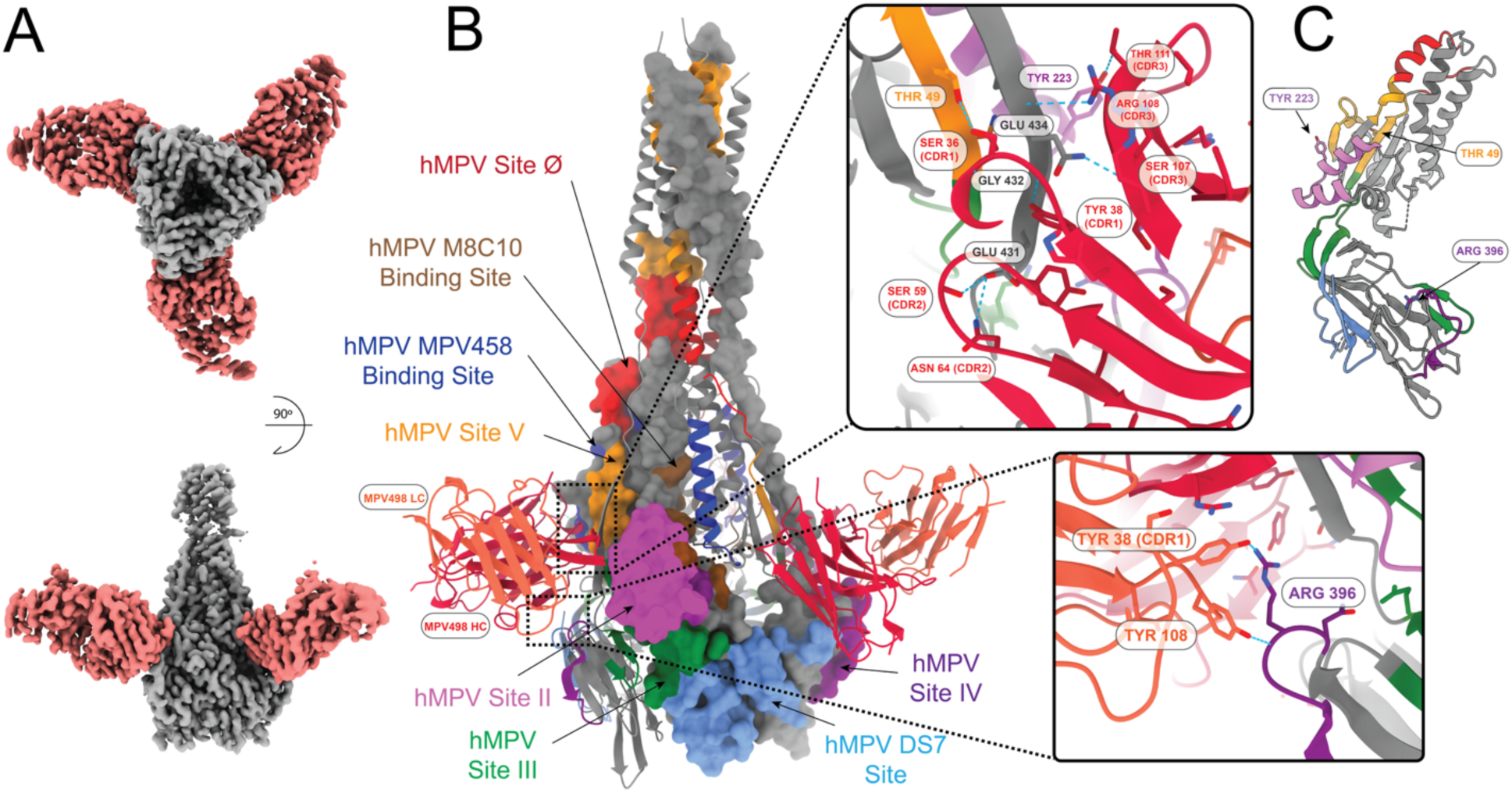
Cryo-EM structure of MPV498 binding hMPV B2 post-fusion F protein. (A) Cryo-EM map for hMPV B2 post-fusion F protein (grey) bound to the MPV498 Fabs (light red). (B) Known binding epitopes of the Fabs to hMPV post-fusion (only one F protein monomer is shown in surface view) with the binding interface of the F protein with MPV498 light chain (tomato red) and heavy chain (dark red). Epitopes are colored based on the amino acid sequences for each epitope. (C) Binding residue location in pre-fusion F shown with side chains (GLU431, GLY432, and GLU434 are not present in the pre-fusion construct of hMPV F DsCavEs2).

As mAbs MPV499 and MPV513 appeared to compete with the intratrimer mAb MPV458, we determined the cryo-EM structures of the Fabs of MPV499 and MPV513 **(Figure 2B)**, each in complex with the hMPV pre-fusion DsCavEs2 F protein **(Figure 4, Figure S2B, Figure S2C, Figure S4, and Figure S5, Table S3)**. Importantly, the hMPV DsCavEs2 F protein construct was shown to form both monomers and trimers^60^. A 6.57 Å resolution map obtained from 22,000 particles for the Fab MPV499-hMPV F DsCavEs2 protein complex **(Figure S4)** showed the Fab of MPV499 binds inside the trimeric interface of the pre-fusion hMPV DsCavEs2 F protein **(Figure 4A)**. For Fab MPV513, a 3.94 Å resolution map was obtained from 208,000 particles **(Figure S5)**, and the binding epitope for Fab MPV513 was also located inside the trimer interface of the hMPV DsCavEs2 F protein, overlapping Fab MPV499 **(Figure 4A-C)**. Given the comparable binding behavior of MPV499 and MPV513, along with the high degree of similarity observed in their low-resolution reconstructions bound to the hMPV DsCavEs2 F protein, we did not pursue extensive data collection for MPV499. It is reasonable to expect that higher-resolution analysis would reveal the same binding mode. The HC of Fab MPV513 interacted via hydrogen bonds through TYR37 (CDR1) and ASP62 (CDR2) with ARG248 and ARG329 residues, respectively, which line the trimeric interface of the hMPV DsCavEs2 F protein **(Figure 4B)**. Additionally, amino acids LEU67 (FR3) and SER74 (FR3) of the LC Fab MPV513 formed hydrogen bonds with ASN202 and GLU70 residues, respectively, which are also located in the trimeric interface **(Figure 4B)**. Two mAbs were previously identified to target the hMPV F trimer interface: MPV458 that targets the amino acids 66-87 at the N-terminal of the F2 fragment^35^, and M8C10 which targets a buried epitope near HRA in the trimeric interface **(Figure 4D)**^36^. Comparing with these two mAbs, three amino acid residues from the binding epitope of Fab MPV513 were shared with the two mAbs: GLU70 targeted by Fab MPV458 and ARG248 and ARG329 targeted by Fab M8C10. Additionally, a fourth unique new amino acid ASN202 located in the trimer interface, was also targeted by Fab MPV513 **(Figure 4B).** Unlike both Fabs MPV458 and M8C10, the unique epitope of Fabs MPV499 and MPV513 is longitudinal and spans from the top to the bottom of the trimeric interface **(Figure 4B)**. Also, the gene usage for these mAbs was variable: IGHV4-31 (MPV499), IGHV5-10 (MPV513), IGHV3-30 (MPV458), and IGHV3-3 (M8C10) for the HC, and IGKV1-39 (for both MPV499 and MPV513), IGKV1-33 (MPV458), and IGKV1-5 (M8C10) for the LC. Additionally, the CDR amino acid lengths were variable for the HC; MPV499 has 10 amino acids at CDR1, 7 at CDR2, and 13 at CDR3, both MPV513 and M8C10 have the same CDR lengths (8.8.13), whereas MPV458 has shorter CDR lengths (8.8.8). In contrast, all four mAbs have the same LC CDR lengths (6.3.9), demonstrating the diversity of genes targeting this epitope.

**Figure 4.**
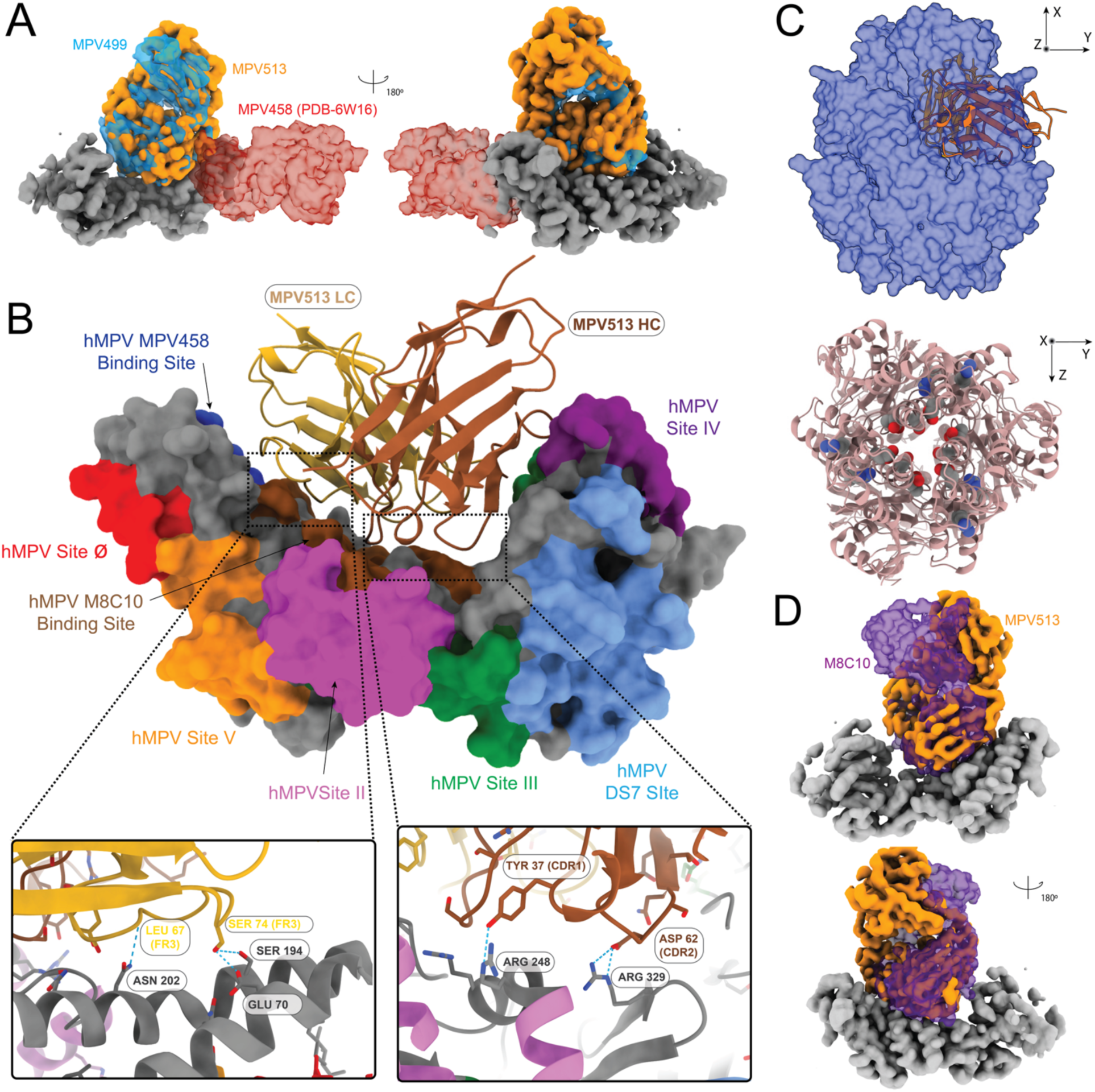
Cryo-EM structure of MPV499 and MPV513 bind hMPV DsCavES2 pre-fusion. **F**. (A) Cryo-EM map for hMPV DsCav-ES2 pre-fusion hMPV protein (grey) bound to the MPV513 Fab (Orange) compared with the MPV499 (transparent blue) and MPV458 (transparent red) from PDB 6W16. (B) Known binding epitopes on hMPV pre-fusion hMPV DsCavEs2 shown in surface view with the binding interface of the F protein with MPV513 light chain (gold) and heavy chain (brown). Epitopes are colored based on the amino acid sequences for each epitope. (C) MPV513 shown in cartoon inside the hMPV DsCav-ES2 pre-fusion F model as a transparent (50%) surface obtained from this study, the Fab is bound to one of the pre-fusion monomers (upper), and binding residues from panel C shown in sphere mode in the hMPV DsCav-ES2 pre-fusion F model from this study, showing their buried position in top view (down). (D) Cryo-EM map for hMPV DsCav-ES2 pre-fusion hMPV protein (grey) bound to the MPV513 Fab (Orange) compared with the M8C10 (transparent purple) from PDB 8T9Z.

Finally, we determined the cryo-EM structure of Fab MPV510 in complex with the hMPV pre-fusion DsCavEs2-IPDS protein, a construct of the hMPV F protein that is stabilized in the pre-fusion conformation and includes interprotomer disulfide bonds that force the hMPV F protein in the closed conformation^51^. A 2.88 Å resolution map was obtained from 287,000 particles **(Figure 5, Figure S2D, Figure S6, Table S3)**. The binding epitope is fully resolved and modeled in 3D **(Figure 5A)**. It revealed that Fab MPV510 binds to an epitope overlapping the epitope previously described by the mAb DS7^65^, and Fab MPV510 interacts with hMPV F through the CDR1 region of the HC and FR2 of the LC **(Figure 5B)**. SER34 (CDR1) of the HC interacted with both ASP331 (located close to the DS7 site) and GLY34 (DS7 site), and ASP36 (CDR1) interacted with LYS20 (DS7 site) **(Figure 5B)**. Additionally, the LC TYR55 (FR2) formed a hydrogen bond with VAL436 (HRB) **(Figure 5B)**. Overlaying the MPV510 structure with the DS7 mAb in complex with hMPV pre-fusion F (PDB 4DAG) revealed an overlapping epitope of both mAbs (**Figure 5C)** with a similar electrostatic binding interface **(Figure 5D and Figure 5E)**. Nevertheless, the encoding genes for both MPV510 and DS7 mAbs were variable: IGHV4-30 and IGKV1-39 for MPV510, while IGHV3-53 and IGLV3-1 for DS7. Also, the CDR amino acid lengths of the HC for both mAbs were variable: 10 (CDR1), 7 (CDR2), and 13 (CDR3) for MPV510, and 8.7.16 for DS7, whereas both mAbs have similar LC CDR lengths (6.3.10 and 6.3.9 for MPV510 and DS7, respectively).

**Figure 5.**
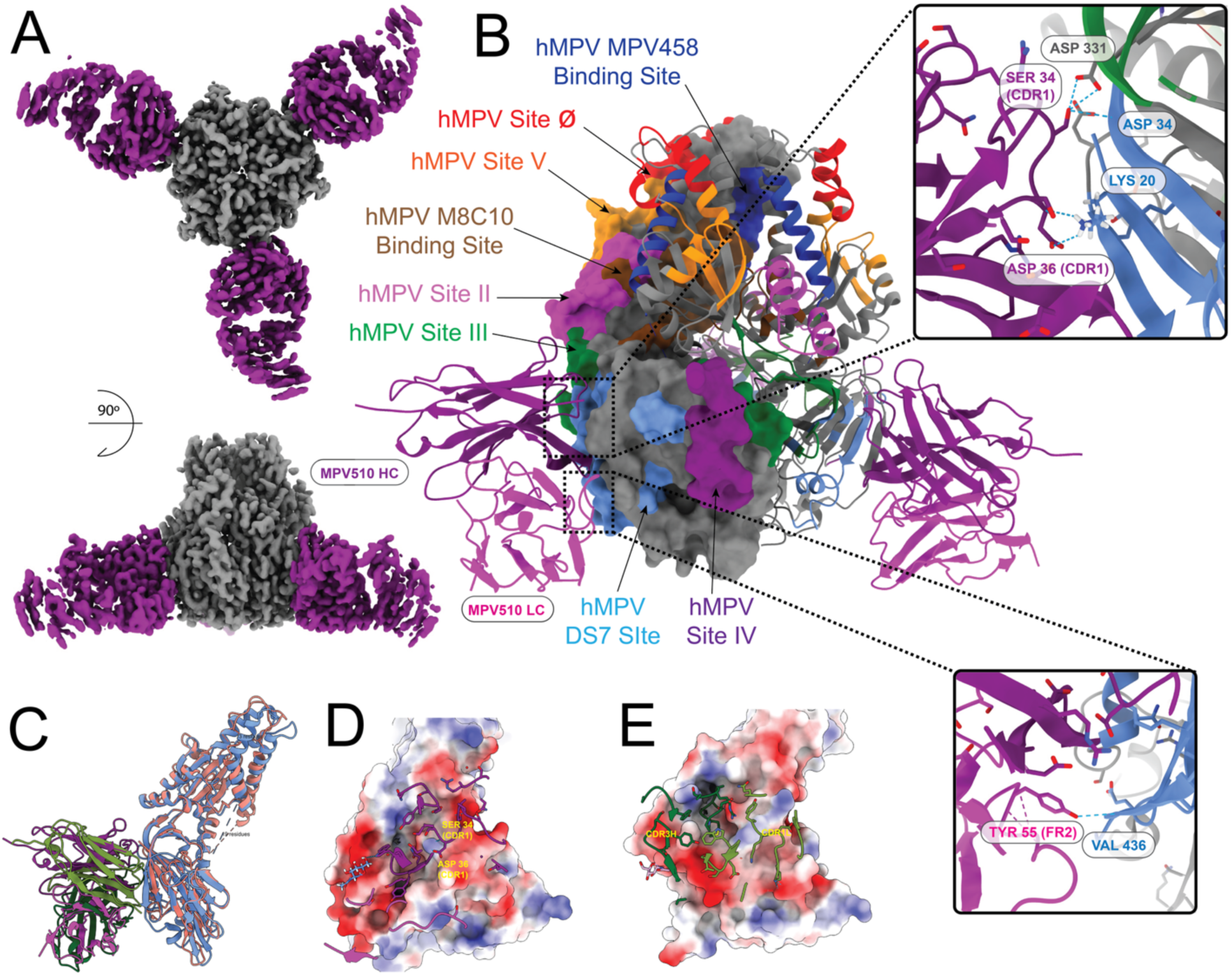
Cryo-EM structure of MPV510 binds to the hMPV DsCavEs2-IPDS F protein. (A) Cryo-EM map for hMPV DsCavEs2-IPDS pre-fusion F protein (grey) bound to the MPV510 Fabs (purple). (B) Known binding epitopes (only one F protein monomer is shown in surface view) with the binding interface of the F protein with MPV510 heavy chain (top) and light chain (bottom). Epitopes are colored based on the amino acid sequences for each epitope. (C) The hMPV F protein monomer shown in salmon-red, MPV510 in purple as previous panels, superimposed with hMPV pre-fusion F-DS7 complex (PDB 4DAG), F shown in blue and the Fab shown in green. (D) Binding interface of MPV510 to DsCav-ES2-IPDS monomer (shown as electrostatic surface). (E) Binding interface of DS7-Fab bound to hMPV F protein (shown as electrostatic surface).

### Prophylactic activity of the isolated mAbs in mice

To determine the prophylactic activity of the five isolated mAbs, 13-week-old female BALB/c mice were treated intraperitoneally with 10 mg/kg of each mAb, positive control MPE8 mAb, or mock-treated with PBS 24 hrs before intranasal infection with hMPV TN/93-32 virus at an infectious dose of 5.0×10^6^ PFU/mouse. At 5 days post-infection, lung virus titers were determined using a plaque assay **(Figure 6A)**. None of the mAb-treated groups with either the mAb MPE8 positive control or the five tested mAbs showed any detectable virus, whereas plaques were detected in the mock-treated group (PBS), indicating the ability of the five isolated mAbs to protect mice against hMPV virus infection **(Figure 6B)**.

**Figure 6.**
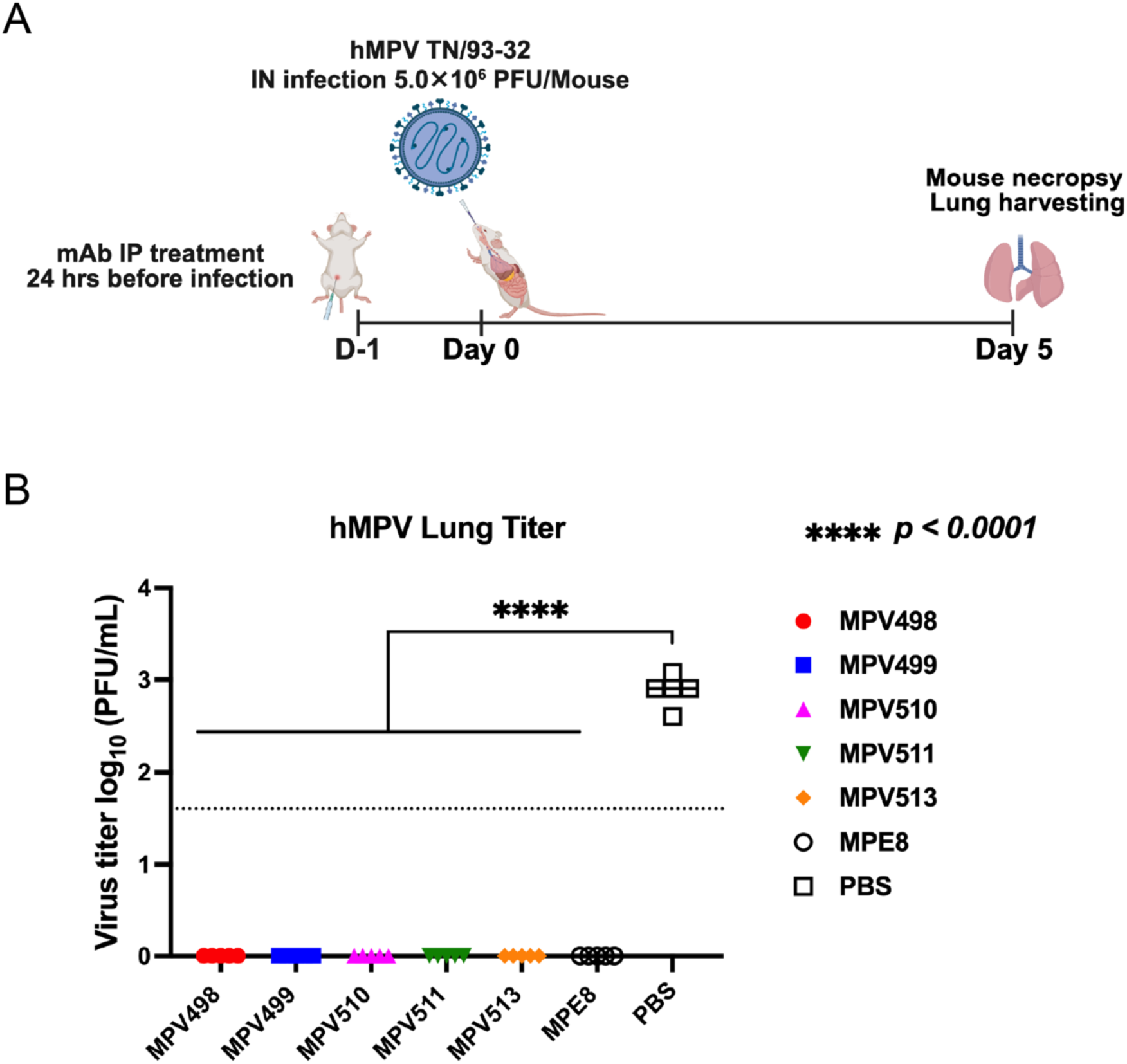
Prophylactic activity of the isolated mAbs against hMPV infection in mice. (A) Experimental plan for testing the prophylactic activity of the mAbs in mice, where 13-week-old female BALB/c mice were treated intraperitoneally with each mAb (10 mg/kg), positive control MPE8 mAb (10 mg/kg), or mock-treated (PBS) 24 hrs before intranasal infection with 5.0X10^6^ PFU/mouse with hMPV TN/93-32. (B) Lung virus titers were determined 5 days post-infection using a plaque assay. The dotted line indicates the limit of detection. * p-value was determined by one-way ANOVA.

## Discussion

Amid the ongoing trials to establish an effective vaccine against hMPV, mAbs are emerging as promising tools for preventing and treating hMPV infections. We and others previously isolated hMPV F-specific mAbs from adults and dissected the antigenic map of the hMPV F protein, whereas the immunological profile of hMPV in children is elusive. In this study, we isolated five mAbs from children previously infected with hMPV. We characterized the repertoire, binding avidity, virus neutralization, *in vivo* prophylactic activity, and structural profiles of these mAbs, which revealed binding to different antigenic sites, including trimeric surface epitopes as well as epitopes located in the trimer interface.

The antibody-encoding gene repertoire for the five isolated mAbs revealed the usage of the VH4 gene family for four mAbs (MPV498, MPV499, MPV510, and MPV511), which was previously reported as one of the main VH gene families encoding several mAbs against hMPV F^20, 30^. Likewise, the IGKV1-39, which comprised the major gene for all isolated mAbs, except mAb MPV511 that utilized V1-5, was also identified as a predominant encoding gene for potent mAbs targeting antigenic sites II and III of the hMPV F^53^. The VH5-10 and VK1-5-encoding genes of mAbs MPV513 and MPV511, respectively, were not commonly reported to generate mAbs targeting hMPV F protein^53^. Intriguingly, our two mAbs targeting the same epitopes at the trimeric interface of hMPV F (MPV499 and MPV513) shared the same DH3-3 and JH4 genes of the HC as well as the VK1-39 gene of the LC, whereas the VH gene was different, where mAb MPV499 utilized VH4-31 while MPV513 utilized VH5-10.

The cryo-EM structure of the Fab MPV498 in complex with the hMPV B2 post-fusion F showed that mAb MPV498 has a unique binding footprint, where the HC grasps the hMPV F trimer along its long axis upward and the LC anchors the antibody to the bottom of the trimer (cradle-like binding) **(Figure 3A and Figure 3B)**. The interactive amino acid residues for the mAb MPV498 epitope were mostly located at the HRB region at the C-terminal end of the F1 subunit, which uncovered a new unidentified epitope at the hMPV post-fusion F (hMPV site VI) **(Figure 3A and Figure 3B)**. Intriguingly, this new epitope corresponds to the newly identified antigenic site VI of the RSV F protein, which is also located at the HRB region at the C-terminal end of the F1 subunit^66^. Since mAb MPV498 binds to both conformations of the hMPV F protein, the putative mechanism of action might be blocking the pre- to post-fusion transition of the F protein when it binds to the pre-fusion F, whereas locking the post-fusion trimer and rendering the F protein fusion-incompetent during the post-fusion binding. Although further structure analysis of the mAb MPV498 with hMPV pre-fusion F is warranted to better understand the mechanism of action of mAb MPV498, the epitope-binning BLI provided some insights into the binding of mAb MPV498 to the hMPV F DsCavEs2 protein. It showed that primary binding of mAb MPV458 to the hMPV F DsCavEs2 protein, which targets an intratrimer alpha helix epitope of amino acids 66-87^35^, completely blocks the access and binding of mAb MPV498 to its epitope, while primary binding of mAb MPV498 then exposing to mAb MPV458 renders partial competition between the two mAbs, indicating that mAb MPV498 might bind to an overlapping epitope distal to the 66-87 helix in the prefusion conformation **(Figure 2B)**. Additionally, the primary binding of mAb MPV498 to the hMPV F DsCavEs2 protein completely blocks the binding of mAb MPV513 (intratrimer-targeting mAb, **Figure 4**) and MPV510 (DS7-targeting mAb, **Figure 5**), implying that the pre-fusion epitope of mAb MPV498 might overlap a region across the trimeric interface and DS7 antigenic sites.

Cryo-EM structures of the MPV499 and MPV513 showed the same binding epitopes on the hMPV DsCavEs2 F protein, where both mAbs target an intratrimer epitope located completely within the hMPV F trimer interface near the middle of the F protein **(Figure 4A)**. Unlike mAb MPV458, where its 66-87 epitope is located only in the C-terminal of the F2 fragment and nearly in the trimeric interface, the epitope of mAbs MPV499 and MPV513 spans the lining interface of the F protein trimeric structure from the C-terminal of F2 to the middle of the F1 subunit **(Figure 4B)**. Interestingly, BLI hMPV F DsCavEs2-coated biosensors showed complete competition when loaded with either mAbs MPV499 or MPV513 and then exposed to mAb MPV458, indicating that the binding of either mAbs MPV499 or MPV513 disrupts the access of mAb MPV458 **(Figure 2B)**. In contrast, loading with mAb MPV458 then exposing to mAbs MPV499 or MPV513 resulted in partial competition, suggesting the ability of mAbs MPV499 or MPV513 to still bind to its epitope, which is located further down in the trimeric interface of the hMPV F protein. While the mechanism of action for the majority of pneumovirus F-targeted mAbs is to hinder the conformational changes from pre-fusion to post-fusion state, mAbs MPV499 and MPV513 might have an alternative way by destabilizing the trimeric structure of the F protein. Also, these findings reinforce the breathing motion of the F trimeric structure previously described for hMPV ^35^ and other virus glycoproteins, including RSV, influenza virus, and HIV^67–72^.

Lastly, we determined the cryo-EM structure for mAb MPV510 in complex with the hMPV pre-fusion DsCavEs2-IPDS at 2.88 Å resolution. It showed that the epitope of mAb MPV510 overlaps with the DS7 antigenic site of the F protein **(Figure 5)**. While mAb MPV510 showed partial competition when exposed to the BLI biosensors loaded with site III-targeted mAbs (MPE8 and MPV364), no competition was observed with DS7-binding MPV196 mAb **(Figure 2B)**. A similar binding profile was noticed with one of our previously identified mAbs, MPV454, which binds to the DS7 site and competes with DS7 mAb, while showing no competition with mAb MPV196^51^. Since the structure of mAb MPV196 has yet to be determined and the epitope of DS7 antigenic site occupies a wide interface, including several amino acid residues^65^, the mAb MPV196 epitope might not completely overlap the DS7 antigenic site, and thus, binding of other mAbs to this site does not block the access of mAb MPV196. Furthermore, the primary binding of mAb MPV510 completely blocks the binding of mAbs MPV458, MPV498, and MPV513, with partial competition with mAb MPV499, implying our previously mentioned finding that binding to the DS7 antigenic site might impact the conformation of the C-terminal region of the hMPV F protein and potentially affecting the accessibility to other antigenic sites or restricting the breathing motion of the trimeric interface that is essential for the access of trimeric interface mAbs **(Figure 2B)**^35, 65^. We also showed that all five mAbs have prophylactic activity in mice in protection against hMPV infection **(Figure 6).** One of the study limitations is to structurally study the binding of mAb MPV498 to hMPV pre-fusion F protein for a better understanding of its mechanism of action.

With the current lack of knowledge of the immunological profile of hMPV infection in children, we isolated five mAbs from children infected with hMPV and provided novel insights into the B-cell repertoire and immunodominant antigenic epitopes of hMPV F protein. Although further studies are necessary to dissect the complete immunological repertoire of hMPV in children, we discovered a new epitope target on the hMPV F protein in children for neutralizing mAbs. Identification of these human mAbs as potential therapeutic tools against hMPV infection is promising, particularly amid the problematic vaccine-enhanced disease observed in association with either recent mRNA or formalin-inactivated hMPV vaccines. Although the generation of human mAbs for clinical use is currently costly, advanced technologies, e.g., the adeno-associated virus (AAV) platform, are rapidly growing for the delivery of mAbs into human cells, providing long-term production of mAbs and subsequently lowering the manufacturing cost^73, 74^.

## Materials and Methods

### Ethics Statement

The mouse studies were approved by the Florida State University Animal Use and Care Committee under protocol IPROTO202300000025. The human subject study was approved by the Institutional Review Board of the University of Michigan Health Sciences and Behavioral Sciences (HSBS IRB) and by the Comité Institucional de Revisión Ética (CIRE) in Nicaragua. Written informed consent was obtained from parents or legal guardians of all participants, and assent was obtained from children ≥6 years of age.

### Blood Collection

Peripheral blood samples were collected from participating children at routine study visits and during acute respiratory illness episodes. Age-appropriate volumes of blood (generally 3-7mL) were obtained by trained study nurses or phlebotomists. Samples were transferred immediately to the local laboratory and processed on-site. Briefly, peripheral blood mononuclear cells (PBMCs) were isolated using density gradient centrifugation. The PBMC layer was collected and washed in cold phosphate-buffered saline (PBS). Cells were then resuspended in a cryopreservation medium [Fetal bovine serum (Corning, product # 35-010-CV) with 10% dimethyl sulfoxide (Sigma, product # 34869), and aliquoted into cryovials at a concentration of 1x10^7^ per ml. Cryovials were gradually frozen using a controlled-rate cooling method (overnight at -80 °C in a Mr. Frosty freezing container) and stored long-term in liquid nitrogen.

### Production and synthesis of recombinant hMPV F protein

Plasmids encoding hMPV A1, A2, B1, B2 F ^51^, pre-fusion stabilized hMPV F DsCavEs2 protein^60^, and hMPV F DsCavEs2-IPDS^51^ recombinant proteins were synthesized (GenScript) and cloned into the pcDNA3.1+ vector. Plasmids were transformed into DH5α *E. coli* (Thermo Scientific) with ampicillin resistance and expanded using the E.N.Z.A. plasmid maxiprep kit (Omega BioTek, Catalog #D6922-04) following the manufacturer’s instructions. Plasmids were then mixed with polyethylenimine (PEI; PolySciences Inc. Catalog #24765) at a 1:4 ratio in OptiMEM cell culture medium (Thermo Scientific, Catalog #31985070) and incubated for 30 min before transfection. hMPV F DsCavEs2 and hMPV F DsCavEs2-IPDS encoding plasmids were co-expressed with human furin (Genscript) at a ratio of 4:1. The DNA-PEI mixture was then transfected into 10^6^ cells/mL Freestyle293F cells (Catalog #R79007) in Freestyle 293 expression medium (Thermo Fisher, Catalog #12338018). After 5 days of incubation, the cultures were centrifuged at 6,000 × g to pellet the cells. The supernatant was filtered through a 0.45 μm sterile filter. The recombinant proteins were purified directly by affinity chromatography using HisTrap Excel columns (Cytiva Life Sciences, Catalog #17371206). Before loading the supernatant onto the column, it was washed with 5 column volumes of wash buffer containing 20 mM Tris HCl (pH 7.5), 500 mM NaCl, and 20 mM imidazole. After passing the supernatant through the column, it was washed with the same wash buffer to reduce nonspecific binding and eluted with buffer containing 20 mM Tris HCl pH 7.5, 500 mM NaCl, and 250 mM Imidazole. After elution, the proteins were concentrated with Amicon Ultra-15 centrifugal units with a molecular cutoff of 30 KDa (Sigma, Catalog #UFC9030).

### Trypsinization and post-fusion induction of hMPV F

To generate homogeneous cleaved trimeric hMPV F, TPCK (L-1-tosylamido-2-phenylethyl chloromethyl ketone)-trypsin (Thermo Scientific, Catalog #20233) was dissolved in double-distilled water (ddH_2_O) at 1 mg/mL. Purified hMPV B2 F protein was incubated with 5 TAME (p-toluene-sulfonyl-L-arginine methyl ester) units/mg of TPCK-trypsin for 1 hr at 37 °C. Trimeric and monomeric hMPV B2 F proteins were purified from the digestion reaction mixture by size exclusion chromatography on a Superdex 200 Increase 10/300 GL (GE Healthcare Life Sciences, Catalog #28990944) in column buffer (50 mM Tris pH 7.5, and 100 mM NaCl). Trimeric hMPV B2 F protein was identified by a shift in the elution profile from monomeric hMPV B2 F protein. The fractions containing the trimers and monomers were concentrated using 30-kDa Spin-X UF concentrators (Corning, Catalog #431489). To obtain fully post-fusion hMPV F, samples were heated at 55 °C for 20 minutes to induce conversion of remaining pre-fusion hMPV F proteins to the post-fusion conformation as previously described^35^.

### Generation of recombinant mAbs

mAb MPV498 was generated by human hybridoma technology as previously described^54^. The mAb V(D)J regions were cloned using RT-PCR and sequenced by Sanger Sequencing, and the mAb was generated recombinantly and cloned into pTwist CMV hIgG1. The rest of the mAbs were isolated by antigen-specific flow sorting and expanded on engineered NIH 3T3 cells that express human CD40L, human interleukin-21 (IL-21), and human BAFF as previously described^57^. Briefly, human PBMCs were washed twice with FACS buffer and then resuspended with 1 mL FACS buffer. The cells were treated with 5% Fc receptor blocker (BioLegend, Catalog #422301) for 30 min and then stained with following antibodies: anti-human CD19-APC (BioLegend, Catalog #302211), anti-human IgM-FITC (BioLegend, Catalog #988306), anti-human IgD-FITC (BioLegend, Catalog #307807), Ghost Dye Red 710 (Tonbo Biosciences, Catalog #13-0871-T500), and streptavidin-PE (Biolegend, Catalog #405203) and streptavidin-BV605-(Biolegend, Catalog #405229) conjugated hMPV B2 F protein, which was biotinylated using the EZ-Link NHS-PEG4-Biotin kit (Thermo Scientific, Catalog #21425). Antigen-specific B cells were gated as CD19^+^/IgM^−^/IgD^−^/Ghost dye^−^/PE^+^/BV605^+^ and sorted one cell per well in 384-well plates pre-coated with gamma irradiated NIH 3T3 cells in StemCell ClonaCell-HY Medium A (StemCell Technologies, Catalog #03801) containing 6.3 μg/mL of CpG (phosphorothioate-modified oligodeoxynucleotide ZOEZOEZZZZZOEEZOEZZZT; Invitrogen) and 1 μg/mL of cyclosporine (MilliporeSigma, Catalog #239835). After 6 days, culture supernatants were screened by ELISA as described below to determine the binding to recombinant hMPV B2 F protein before culturing for another 24 hours at 37°C. The following day, the wells with positive binding were selected for RNA extraction using the Qiagen RNeasy Micro kit (Qiagen, Catalog #74004). Cells were gently scraped from the bottom of the well before proceeding with the manufacturer’s RNA extraction protocol. RNA samples were stored at -80C until they were reverse transcribed to cDNA with the SuperScript IV Synthesis System (ThermoFisher, Catalog #18091050). The variable region sequences of immunoglobulin G (IgG) heavy/light chains were determined by nested PCR as previously described^58^. Based on the usage of V/D/J gene alleles, cloning PCR primers were picked for cloning PCR with the first PCR products as the template. Purified cloning PCR products of heavy/light chains were cloned into expression vectors (AbVec-hIgG1 and AbVec-hIgKappa expression vectors), and the plasmids were sent to Genewiz for sequencing. After confirming all the sequences are correct, the heavy chain/light chain (HC/LC) plasmids were transformed into DH5α, and plasmids were prepared using the E.Z.N.A. Plasmid Maxi Kit (Omega Bio-Tek, Catalog #D6922-04). Recombinant mAbs were expressed by transfecting Expi293F cells (ThermoFisher, Catalog #A14527) using the Freestyle293 transfection protocol described above, or transfecting ExpiCHO-S cells (ThermoFisher, Catalog #A29127) using the ExpiFectamine CHO transfection kit (ThermoFisher, Catalog #A29129) with HC/LC plasmids and purified from culture supernatant with Protein G column (Cytiva, Catalog #17040501)^50^.

### ELISA for binding to hMPV F proteins

384-well plates (Greiner Bio-One, Catalog #781162) used for ELISA were coated with 2 μg/mL of the recombinant proteins in PBS and incubated overnight at 4 °C. The plates were then washed once with water, followed by blocking with block buffer comprising 2% nonfat milk (Biorad, Catalog #1706404XTU) supplemented with 2% goat serum (Thermo Fisher, Catalog #16210072) in PBS for 1 hr at 37 °C. The plates were washed again three times with water. A total of 25 μL of the serially diluted primary antibodies with a starting concentration of 20 μg/mL were added to the wells and incubated for 1 hr at 37 °C, followed by three washes with water. Goat anti-human IgG Fc AP secondary antibody (Southern Biotech, Catalog #2048-04) diluted in block buffer (1:4,000) was next applied to the wells and incubated again at 37 °C for 1 hr. The plates were washed with PBS and 0.05% Tween 20 (PBS-T) three times, and 25 μL of p-nitrophenyl phosphate diluted to a concentration of 1 mg/mL in a buffer containing 1M Tris base and 0.5 mM magnesium chloride, having a pH of 9.8, was added. Plates were then read at the absorbance of 405 nm on a BioTek plate reader. The binding assay data were analyzed on GraphPad Prism using a nonlinear regression curve fit and log(agonist)- versus-response function for calculating the EC_50_ values.

### Virus Plaque Reduction Neutralization Assay

One day before the assay, LLC-MK2 cells in Opti-MEM (Thermo Scientific, Catalog #M352-500) supplemented with 2% fetal bovine serum (Corning, product # 35-010-CV) and 1% antibiotic-antimycotic (Gibco, Catalog #15240062) were plated on 24-well plates (10^5^ cells /well) and incubated at 37°C in a CO_2_ incubator. The following day, three-fold serially diluted mAbs were then mixed with each hMPV virus strain (CAN/97-83 or TN/93-32) in equal amounts (1:1) in Opti-MEM and incubated for 1 hr at room temperature. The virus-antibody mixture was added onto the LLC-MK2 cells after washing with PBS three times. The mixture was incubated at 37°C for 1 hr with constant rocking. The cells were next overlaid with 0.75% methylcellulose dissolved in Opti-MEM I supplemented with 5 μg/mL of trypsin-EDTA (Thermo Scientific, Catalog #25200056) and 100 μg/mL of CaCl_2_. The cells were incubated for 5 days. Cells were then fixed with 4% neutral buffered formalin (Epredia, Catalog #22-110-869) and blocked with block buffer comprising 2% nonfat milk (Biorad) supplemented with 2% goat serum in PBS for 1 hr at 37°C. Then, the plates were washed three times with water, and mAb MPV364 (primary mAb) was added to a final concentration of 1 μg/mL (1:1,000 dilution) in 1% blocking solution for 1 hr at 37°C. The plates were then washed three times with water, and goat anti-human IgG HRP secondary antibody (Southern Biotech, Catalog #2048-05) diluted to a ratio of 1:2,000 in block buffer was added and incubated for 1 hr at 37°C. Plates were washed again with water five times, and 200 μL of TrueBlue peroxidase substrate (SeraCare, Catalog #5510-0030) was added to each well. The plates were incubated for 5–10 min until the plaques were visible, then the plate was washed once with water. Plaques were counted manually and compared to the virus-only control (100% virus infection well). Briefly, the average of three replicates for the tested antibody-diluted wells was divided by the average of the virus-only control triplicate wells and multiplied by 100. GraphPad Prism was used to calculate the IC_50_ values using a nonlinear regression curve fit and the log(inhibitor)-versus-normalized response function.

### Epitope binning using biolayer interferometry

An initial baseline in running buffer (PBS, 0.5% bovine serum albumin [BSA], and 0.05% Tween 20) for 60s was obtained first before loading the anti-Histidine (Anti-His) biosensor tips (Gator Bio, Catalog #160009) with 50 μg/ml of His-tagged hMPV DS-CavEs2 F protein for 120 s. The baseline signal was measured again for 60 s before biosensor tips were immersed into wells containing 50 μg/ml primary antibody for 300 s. Following this, biosensors were immersed into wells containing 50 μg/ml of a second mAb for 300 s. Percent binding of the second mAb in the presence of the first mAb was determined by comparing the maximal signal of the second mAb after the first mAb was added to the maximum signal of the second mAb alone. mAbs were considered noncompeting if the maximum binding of the second mAb was ≥67% of its uncompetitive binding. A level of between 34% and 66% of its uncompetitive binding was considered partial competition, and ≤33% was considered competition. For the affinity experiment, hMPV DsCavEs2 F protein (100 µg/mL) were loaded onto anti-His biosensors, followed by exposure to 2-fold serially diluted mAb MPV498 (starting with 100 µg/mL) association (300 s), then dissociation (600 s). BLI Gator data analysis software (Version 2.16.6.0130) was used to analyze the data and calculate the K_on_ values. A reference curve containing buffer only (without MPV498 but with hMPV DsCavEs2 F protein still loaded) was subtracted from each value.

### Cryo-EM sample preparation and data collection

To obtain Fab fragments, papain digestion was performed using the Pierce Fab preparation kit (Thermo Fisher Scientific, Catalog # 44985) according to the manufacturer’s protocol. Fab fragments were purified by running on the Superdex 200 Increase 10/300 GL (GE Healthcare Life Sciences, Catalog #28990944) column according to the manufacturer’s protocol. hMPV F proteins (hMPV B2 post-fusion F, hMPV DsCavES2 pre-fusion F or hMPV DsCavsES2-IPDS pre-fusion F) were purified as mentioned above and incubated with a 3-fold molar excess of the Fabs from each mAb for 60 minutes at room temperature, followed by incubation on ice. The complex was then loaded onto an AKTA FPLC system equipped with a Superdex 200 Increase 10/300 GL (GE Healthcare Life Sciences, Catalog #28990944). Peak fractions were analyzed by SDS-PAGE, and the complex was identified and extracted. Prior to grid preparation, the sample was diluted using 20 mM Tris pH 8.0, 100 mM NaCl. Cryo-EM grid preparation involved applying 4 µL of the sample to a Quantifoil 1.2/1.3 Cu 300 mesh grids (EMS, Q350CR1.3), which had been plasma-cleaned for 20s using a Solarus plasma cleaner (Gatan). Grids were plunge-frozen using a Vitrobot Mark IV (Thermo Fisher) at 10 °C and 100% humidity, with 1.5 s blot time and −1 blot force, then plunged into liquid ethane cooled by liquid nitrogen. After clipping and screening, data were collected on a Titan Krios equipped with a DE-Apollo direct electron detector. A single grid yielded 7,900 (MPV498/B2 post-fusion F), 400 (MPV499/ DsCavEs2 pre-fusion F), 8,000 (MPV510/ DsCavEs2-IPDS pre-fusion F) or 6,100 (MPV513/ DsCavEs2 pre-fusion F) multi-frame micrographs. Images were recorded at a magnification of 59,000X for all mAbs except that of MPV513/hMPV F DsCavEs2 pre-fusion F that was at 105,000X, corresponding to a calibrated pixel size of 0.79 Å/pixel, with a total electron dose of 60 e⁻/Å². Data collection statistics are summarized in **Table S3**.

### Cryo-EM Data Processing

Movie frame alignment and CTF estimation were done using a combination of MotionCor^75^, CTFFIND4^76^, and patch motion correction and patch CTF estimation from the cryoSPARC^77^. All other processing steps were performed in cryoSPARC; initial preprocessing included patch motion correction and CTF estimation, followed by blob-based particle picking. After 3–4 rounds of 2D classification, well-resolved classes were selected, and three templates were generated for template-based particle picking. Particles were extracted with 2x binning (Fourier crop ½) and subjected to ab initio reconstruction to generate 10 distinct models. These served as references for multiple rounds (4–5) of heterogeneous refinement, progressively removing poorly aligned particles. At each step, only particles contributing to well-resolved classes were retained. The final particle stack was re-extracted without binning. One to two additional rounds of heterogeneous refinement were performed to further clean the dataset. The best ab initio model was used for non-uniform refinement, with C3 symmetry applied where appropriate **(see Table S3)**. To mitigate orientation bias, the “Rebalance Orientation” job was used, followed by another round of non-uniform refinement with per-particle CTF refinement and defocus optimization enabled. Final maps were post-processed and sharpened using DeepEMhancer^78^ with the default “tight” model for Figure preparation and visualization.

### Model Building and Refinement

Fab models were generated using AlphaFold3 based on their amino acid sequences^79^. For the hMPV F proteins, existing structures were used as templates: 7M0I for B2 post-fusion F, 6W16 for DsCavEs2 pre-fusion F, and 7UR4 for DsCavEs2-IPDS pre-fusion F. Initial fitting of F protein and Fab models into cryo-EM density maps was performed using UCSF Chimera’s Fit in Map tool^80^. Model refinement was conducted iteratively using PHENIX real-space refinement^81^ with manual adjustments in Coot^82^. All refinements were performed against cryoSPARC-sharpened maps. The final models were evaluated for geometry and map fit using standard validation metrics **(Table S3)**.

### Data Deposition

Atomic coordinates were deposited in the Protein Data Bank (PDB) and the corresponding density maps in the Electron Microscopy Data Bank (EMDB) as described in **Table S3**. Motion-corrected micrographs were deposited in EMPIAR.

### Mouse experiment

To determine the prophylactic efficacy of the five isolated mAbs, 13-week-old female BALB/c mice were treated intraperitoneally with each mAb (10 mg/kg), positive control MPE8 mAb (10 mg/kg), or mock-treated (PBS) 24 hrs before intranasal infection. Mice were infected with 5.0×10^6^ PFU/mouse of hMPV strain TN/93-32 after anaesthetizing mice with 5% isoflurane (Pivetal, NDC #46066-0115-04) using a V-10 mobile laboratory animal anesthesia system (VetEquip, Catalog #901807). Five days following infection, mice were euthanized via Euthasol (Virbac Animal Health, Catalog #51311-0050-01), followed by cervical dislocation, and the lungs were collected. Lungs were harvested, homogenized using a gentleMACS dissociator in M tubes (Miltenyi Biotec, Catalog #130-093-236), and viral titers were determined by plaque immunostaining as previously described^30^. Data were analyzed in GraphPad Prism using the ordinary one-way analysis of variance (ANOVA) function to calculate *P* values.

