## Supplemental Figures and Tables for "Structural Basis for Childhood Antibody Recognition of The Human Metapneumovirus Fusion Protein"

**Table S1. Metadata for the monoclonal antibodies isolated from hMPV-infected children.**

| mAb | Participant ID | Participant Sex | Participant age at infection (year) | Repeated infection (Yes/No) * |
|---|---|---|---|---|
| MPV498 | 9131 | Male | 1 | No |
| MPV499 | 8749 | Male | 1 | No |
| MPV510, MPV511 and MPV513 | 8631 | Female | 1 | No |

*This refers to whether the infection is repeated during the cohort (2011 to 2016).

| Table S2. Monoclonal antibody sequences. | | | | |
|---|---|---|---|---|
| **mAb** | **HC amino acids** | **HC Junction** | **LC amino acids** | **LC Junction** |
| MPV498 | QVQLQESGPGLVKPSETLSLTCTVSGGSISSYYWSWIRQPAGKGLEWIGRIYTSGNTNYNPSLKSRVTMSLDTSKNQVSLKLSSVTAADTAVYYCARSRVATTPVGLRDWLDPWGQGTLVTVSS | CARSRVATTPGLRDWLDPW | TPSLSASVGDRVTITCRASQSISSYLNWYQQKPGKAPKLLIYAASSLQSGVPSRFSGSGSGTDFTLTISSLQPEDFATYYCQQSYSTPLFFGQGTKLEIK | CQQSYSTPLFF |
| MPV499 | QVQLQESGPGLVKPSQTLSLTCTVSGGSISSGDYNWNWIRQHAGKGLEWIGYINYSGSTDYNPSLKSRVTISVDTPKNQFSLKLTSVTAADTAVYYCARGVDFWSGYCDYWGQGSLVTVSS | CARGVDFWSGYCDYW | PVSLSASVGDRVTITCRASQSISTYLNWYQQKPGKAPKLLIYAASSLQSGVPSRFSGSVSGTDFTLTISSLQPEDFATYYCQQTYTTPLTFGGGTKVEIK | CQQTYTTPLTF |
| MPV510 | QVQLQESGSGLVKPSQTLSLTCAVSGGSISSGDSSWSWIRQPPGKGLEWIGHVYESGNTYYDPSLQSRVTISVDRSRNQFSLKLTSVTVADTAVYYCAREGNYGWDYFDYWGQGTLVTVSS | CAREGNYGWDYFDYW | DIQMTXSPFSLSASVGDRVTITCRASQSINSYLNWYQQKPGKAPRLLIYAASSLQSGVPSRFRGSGSGTDFALTISSLQPEDFATYYCQQSYRPPSRTFGQGTKVEMK | CQQSYRPPSRTF |
| MPV511 | QVQLQQWGAGLLKPSETLSLTCGVDGGSFNGYYWNWVRQLPGKGLEWIGEVSYAGSDNYNPSLKSRATISGDRSRKQFSLRLDSVTVAGTGVYYCARDKGVLDYSFGLDVWGQGTTVTVSS | CARGKGVLDYSFGLDVW | DIQMTQSPSTLSASVGDRVTITCRASQSVSNWLAWYQQKSGKAPKLLIYKASSLESGVPSRFSGSGSGTEFTLTISGLQPDDFATYYCQEHNSDSRAFGQGTKVEIK | CQEHNSDSRAF |
| MPV513 | EVQLVQSGAEVKKPGESLRISCKGSGYDFPSYWISWVRQMPGKGLEWMGRIDPTDSNTNYSPSFQGHVTLSADKSISTAYLQWSSLKASDTAIYYCARHSDSWSYYEDSWGQGTLVTVSS | CARHSDFWSYYEDSW | DIQMTQSPSSLSASVGDRVTITCRASQSISTYLNWYQQKPGKAPKLLIYAASSLQSGVPSRFSGRLSGTDFTLTISSLQPEDFATYYCQQTYSSPRTFGQGTKVEIK | CQQTYSSPRTF |

| Table S3. Cryo-EM data collection, refinement, validation, and model-building statistics. | | | | |
|---|---|---|---|---|
| | MPV498/hMPV B2-post fusion F | MPV499/hMPV DsCav-ES2 pre-fusion F | MPV510/hMPV DsCav-ES2-IPDS pre-fusion F | MPV513/hMPV DsCav-ES2 pre-fusion F |
| **Data collection and processing** | | | | |
| Scope | Titan Krios | Titan Krios | Titan Krios | Titan Krios |
| Magnification | 59,000 | 59,000 | 59,000 | 105,000 |
| Voltage (kV) | 300 | 300 | 300 | 300 |
| Electron exposure (e–/Å$^2$) | 60 | 60 | 60 | 60 |
| Defocus range (μm) | 0.8-2.8 | 0.8-2.8 | 0.8-2.8 | 0.8-2.8 |
| Camera | DE-Apollo | DE-Apollo | DE-Apollo | Gatan K3 |
| Pixel size (Å) | 0.79 | 0.79 | 0.79 | 0.88 |
| Symmetry imposed | C3 | C1 | C3 | C1 |
| Initial particle images (no.) | 2,500,000 | 150,000 | 1,200,000 | 1,800,000 |
| Final particle images (no.) | 111,000 | 22,000 | 287,000 | 208,000 |
| Map resolution (Å)<br>FSC threshold | 2.61<br>0.143 | 6.57<br>0.143 | 2.88<br>0.143 | 3.94<br>0.143 |
| Map resolution range (Å) | 2-6 | 5.5-10 | 2.3-7 | 3.5-6 |
| **Refinement** | | | | |
| Initial model used (PDB code) | 7M0I/Alphafold | N/A | 7UR4/Alphafold | 6W16/Alphafold |
| Model resolution (Å)<br>FSC threshold | 2.61<br>0.143 | N/A | 2.88<br>0.143 | 3.94<br>0.143 |
| Model resolution range (Å) | 2-6 | N/A | 2.3-7 | 3.5-6 |
| Map sharpening *B* factor (Å$^2$) | 87.5 | 452 | 105.2 | 234.9 |
| R.m.s. deviations<br>Bond lengths (Å)<br>Bond angles (°) | <br>0.002<br>0.521 | N/A | <br>0.004<br>0.680 | <br>0.003<br>0.602 |
| Validation<br>Clash-score<br>Poor rotamers (%) | <br>6.12<br>2.83 | N/A | <br>24.21<br>3.37 | <br>6.49<br>3.29 |
| Ramachandran plot<br>Favored (%)<br>Allowed (%)<br>Disallowed (%) | <br>96.55<br>3.45<br>0.00 | N/A | <br>92.78<br>6.14<br>1.08 | <br>91.53<br>7.97<br>0.50 |
| PDB accession code | 9OS5 | N/A | 9PDY | 9PDX |
| EMDB accession code | EMD-70793 | EMD-72382 | EMD-71548 | EMD-71547 |
| EMPIAR accession code | TBD | TBD | TBD | TBD |

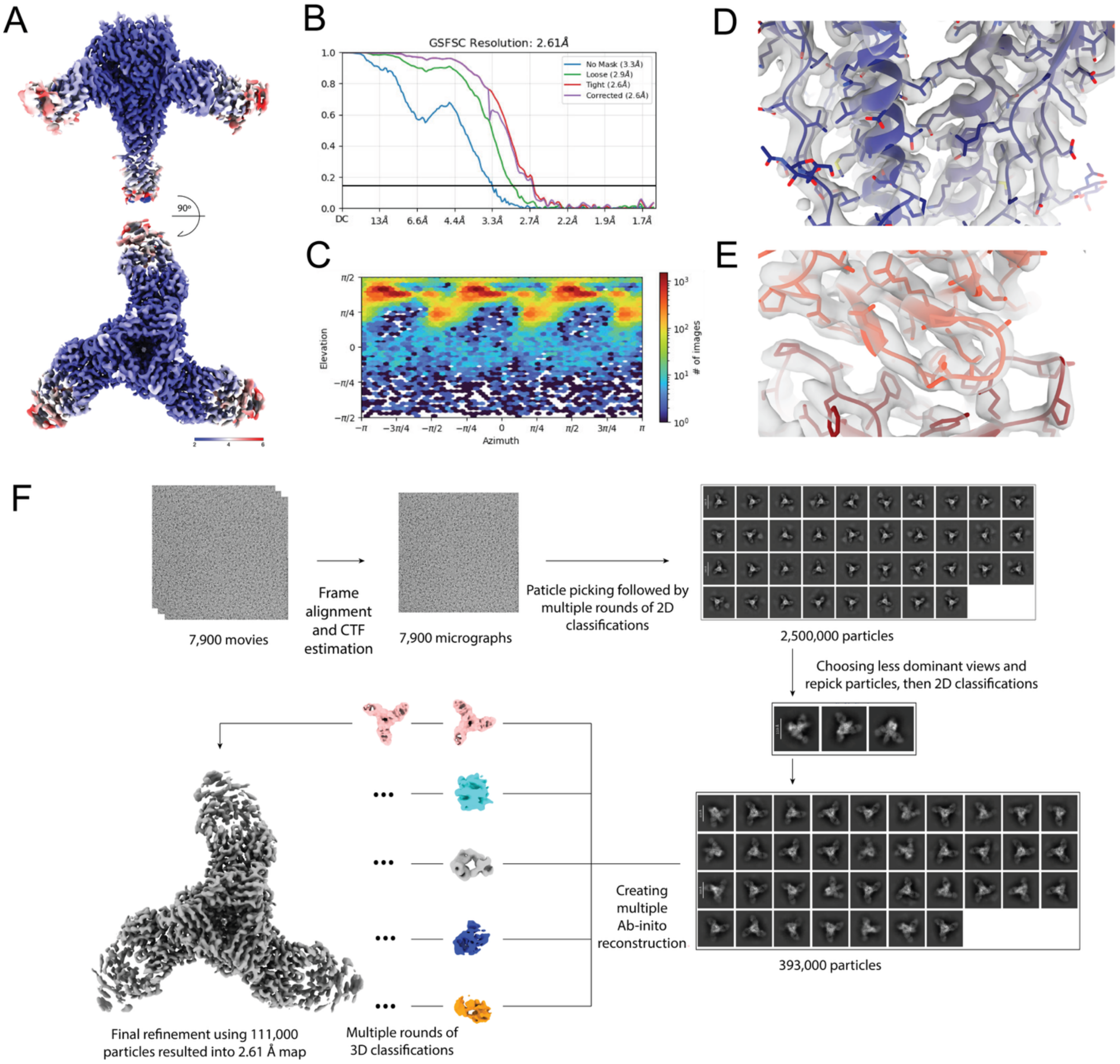


**Figure S1. MPV498 cryo-EM processing workflow.** (A) Local resolution map obtained for hMPV B2 post-fusion F protein bound to the MPV498 Fab. (B) GSFSC curve of the refined Cryo-EM map. (C) Particle distribution map for the final refinement. (D) Model to map fit example of the hMPV B2 post-fusion F protein. (E) Model to map fit example of the MPV498 Fab near the binding site. (F) Overall Cryo-EM data processing workflow.

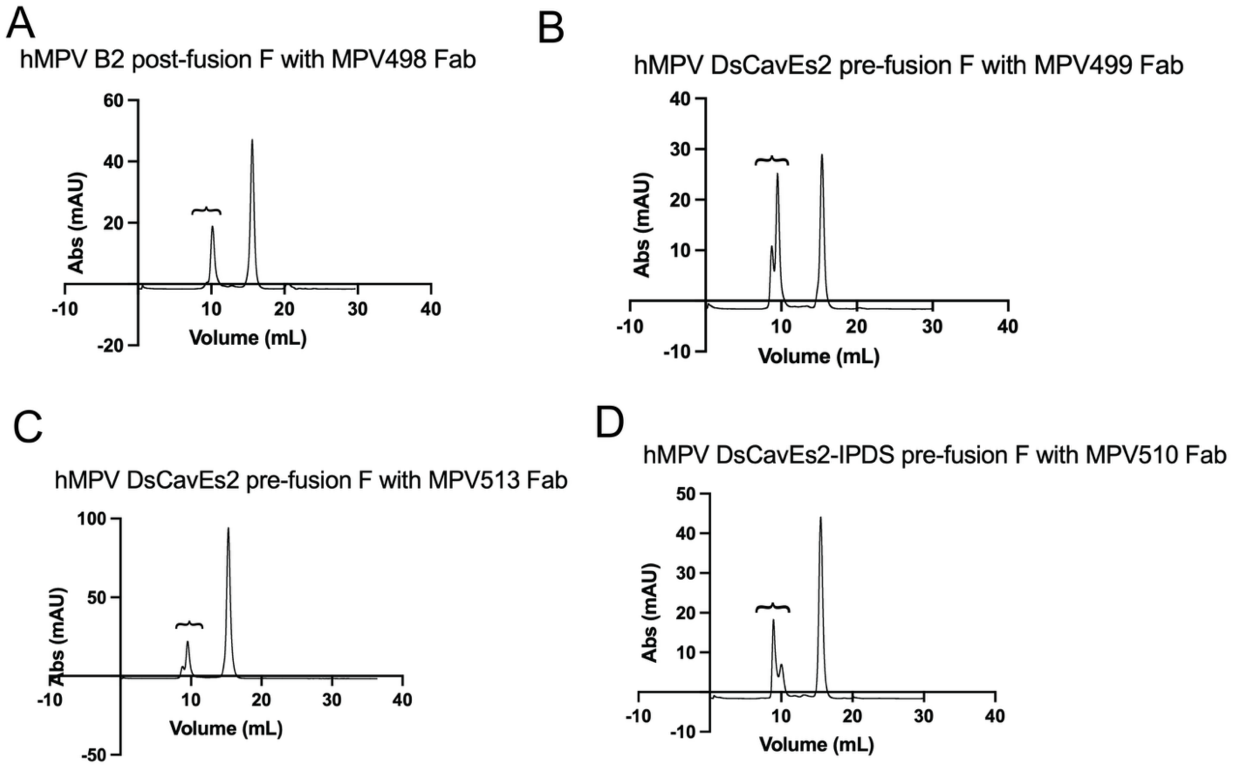


**Figure S2. Size-exclusion curves obtained from hMPV F-Fab complexes.** (A) hMPV B2 post-fusion F with MPV498 Fab. (B) hMPV DsCavEs2 pre-fusion F with MPV499 Fab. (C) hMPV DsCavEs2 pre-fusion F with MPV513 Fab. (D) hMPV DsCavEs2-IPDS pre-fusion F with MPV510 Fab. Selected complex peaks used for grid preparation are shown with accolades.

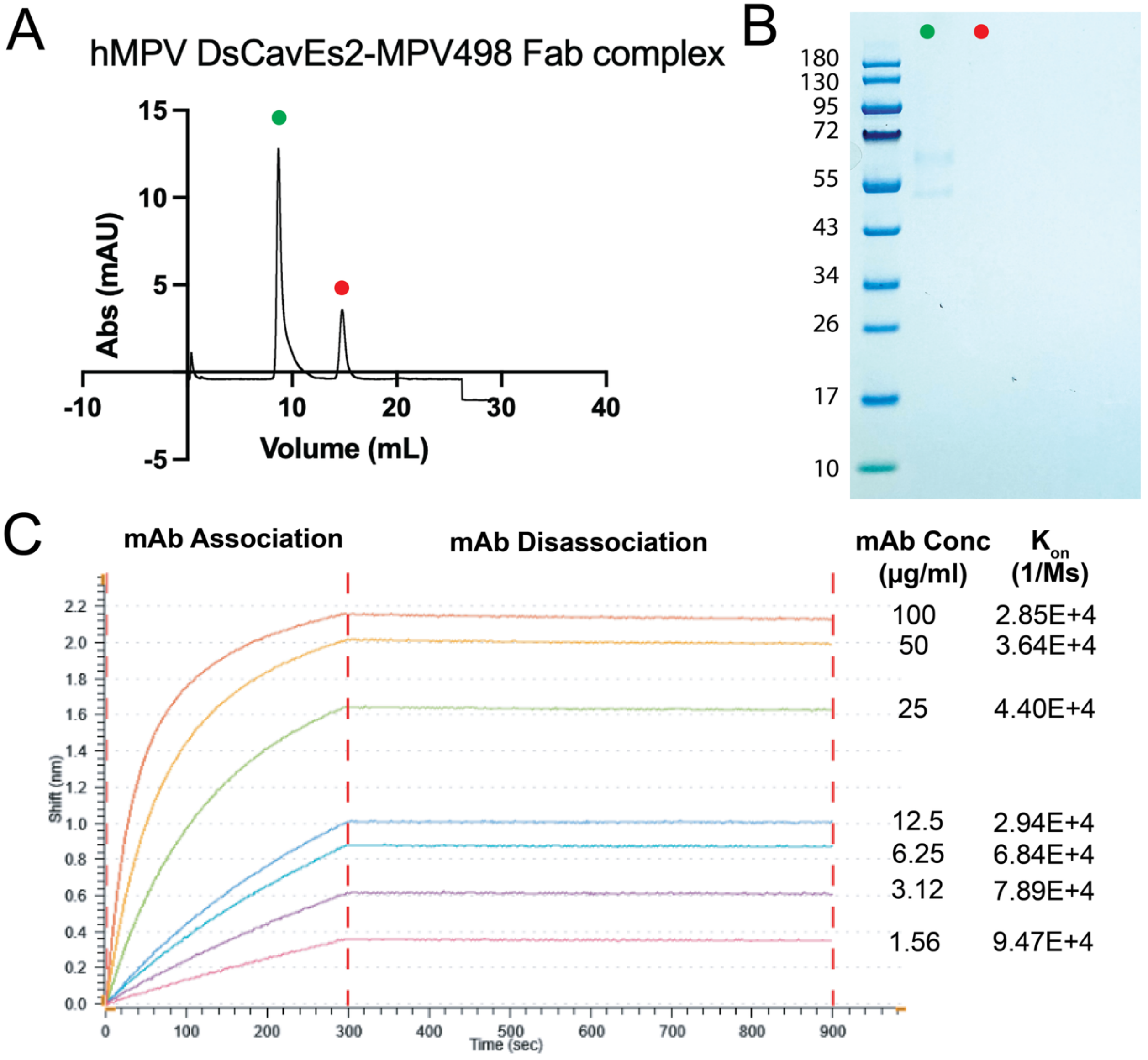


**Figure S3. Binding of MPV498 to hMPV DsCavEs2 F.** (A) Size exclusion curves obtained from AKTA-pure for hMPV DsCavEs2 F-MPV498 Fab complex (green) and extra Fab (red). (B) SDS-PAGE for the hMPV DsCavEs2 F-MPV498 Fab complex (green) and extra Fab (red). (C) Binding affinity of mAb MPV498 to DsCavEs2 F using BLI. hMPV DsCavEs2 protein (100 µg/ml) was loaded onto anti-His biosensors. Processed curves for the associations (300 s) and disassociations (600 s) of 2-fold serially diluted mAb MPV498 (starting with 100 µg/ml) are shown. $K_{on}$ values are listed for each mAb MPV498 dilution. A reference curve containing buffer only was subtracted from each value.

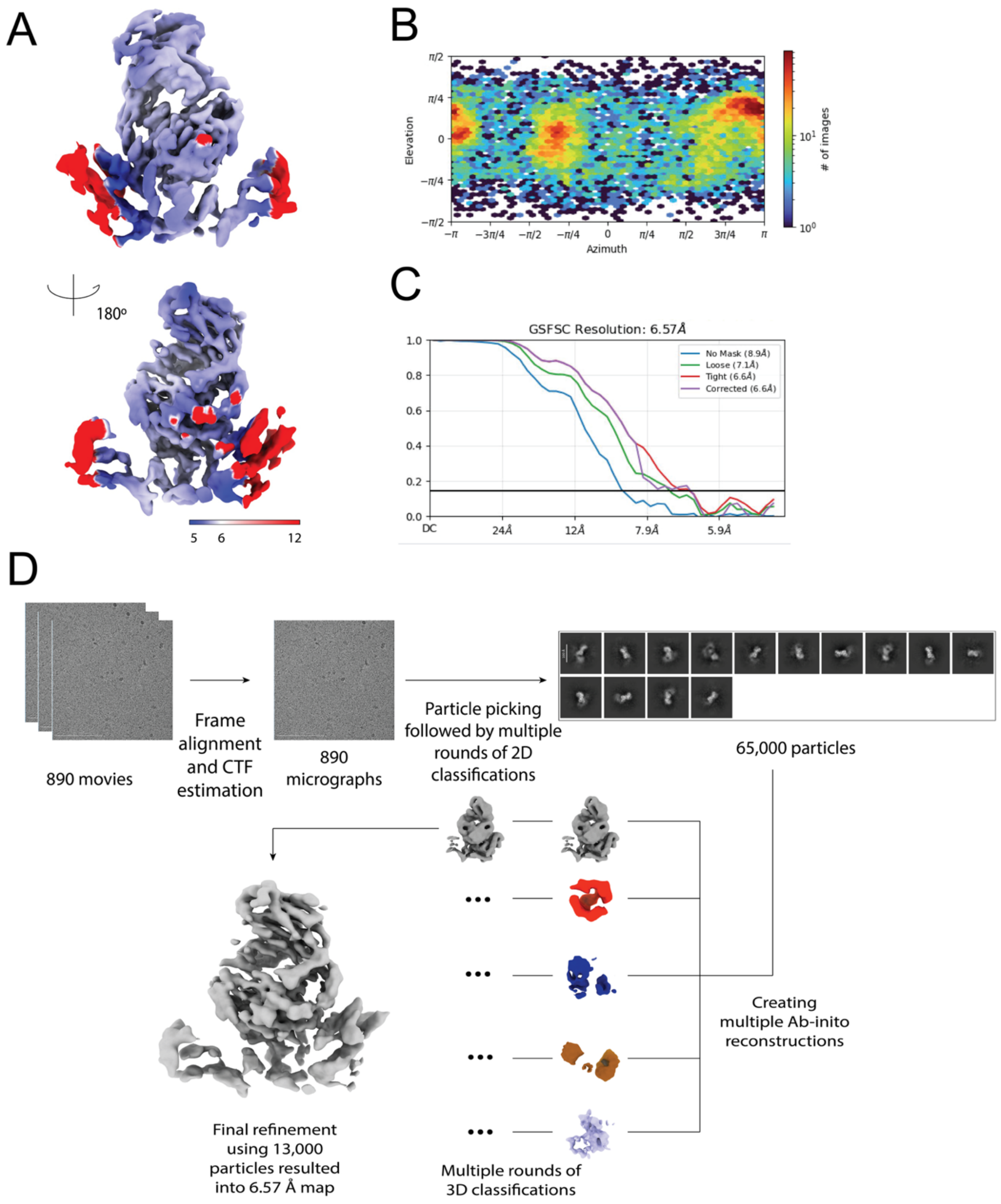


**Figure S4. MPV499 cryo-EM workflow**. (A) Local resolution map obtained for hMPV DsCav-ES2 pre-fusion F protein bound to the MPV499 Fab. (B) GSFSC curve of the refined Cryo-EM map. (C) Particle distribution map for the final refinement. (D) Overall Cryo-EM data processing workflow.

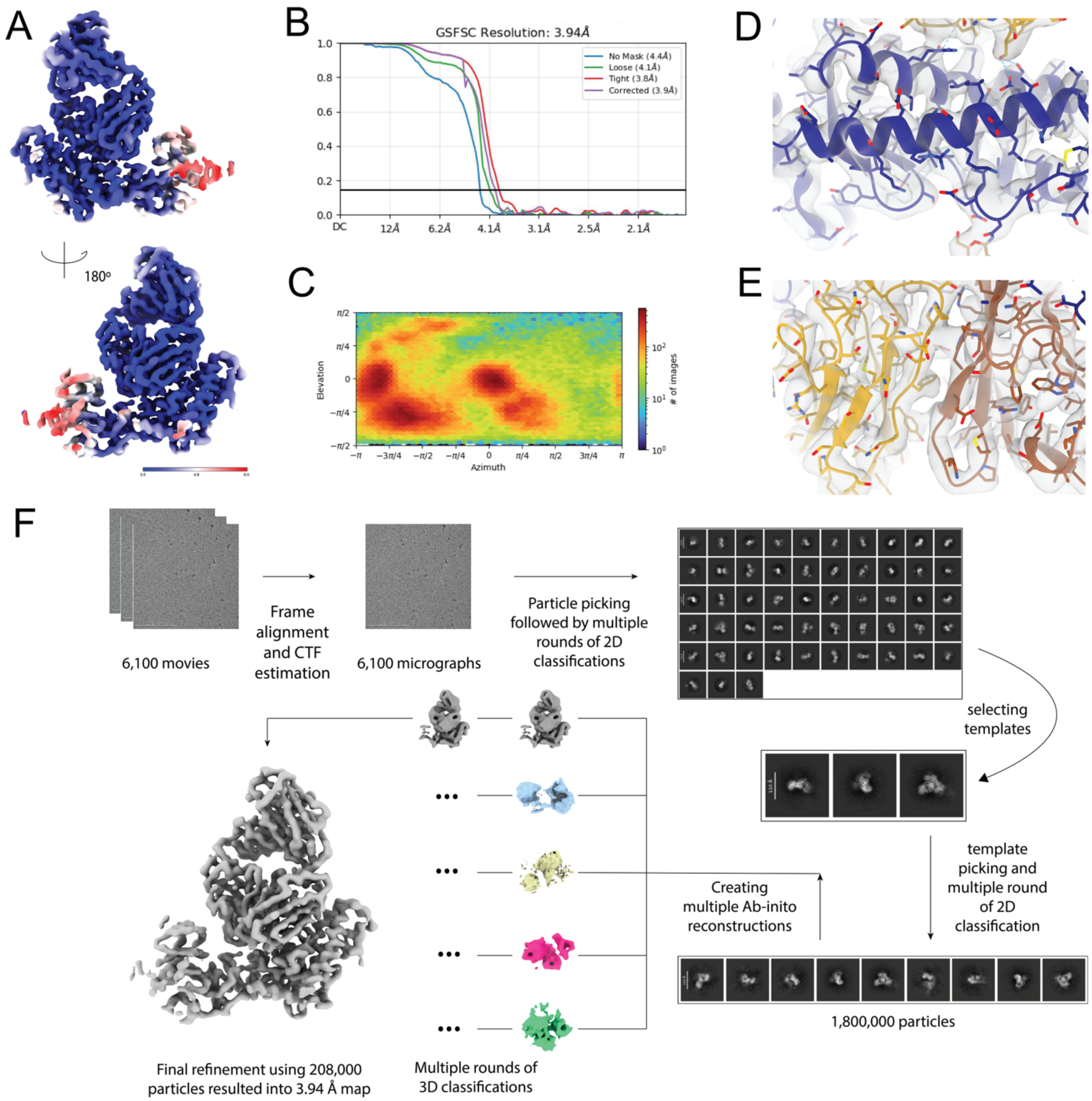


**Figure S5. MPV513 cryo-EM workflow**. (A) Local resolution map obtained for hMPV DsCav-ES2 pre-fusion F protein bound to the MPV513 Fab. (B) GSFSC curve of the refined Cryo-EM map. (C) Particle distribution map for the final refinement. D) Model to map fit example of the hMPV DsCav-ES2 pre-fusion F protein. (E) Model to map fit example of the MPV513 Fab. (F) Overall Cryo-EM data processing workflow.

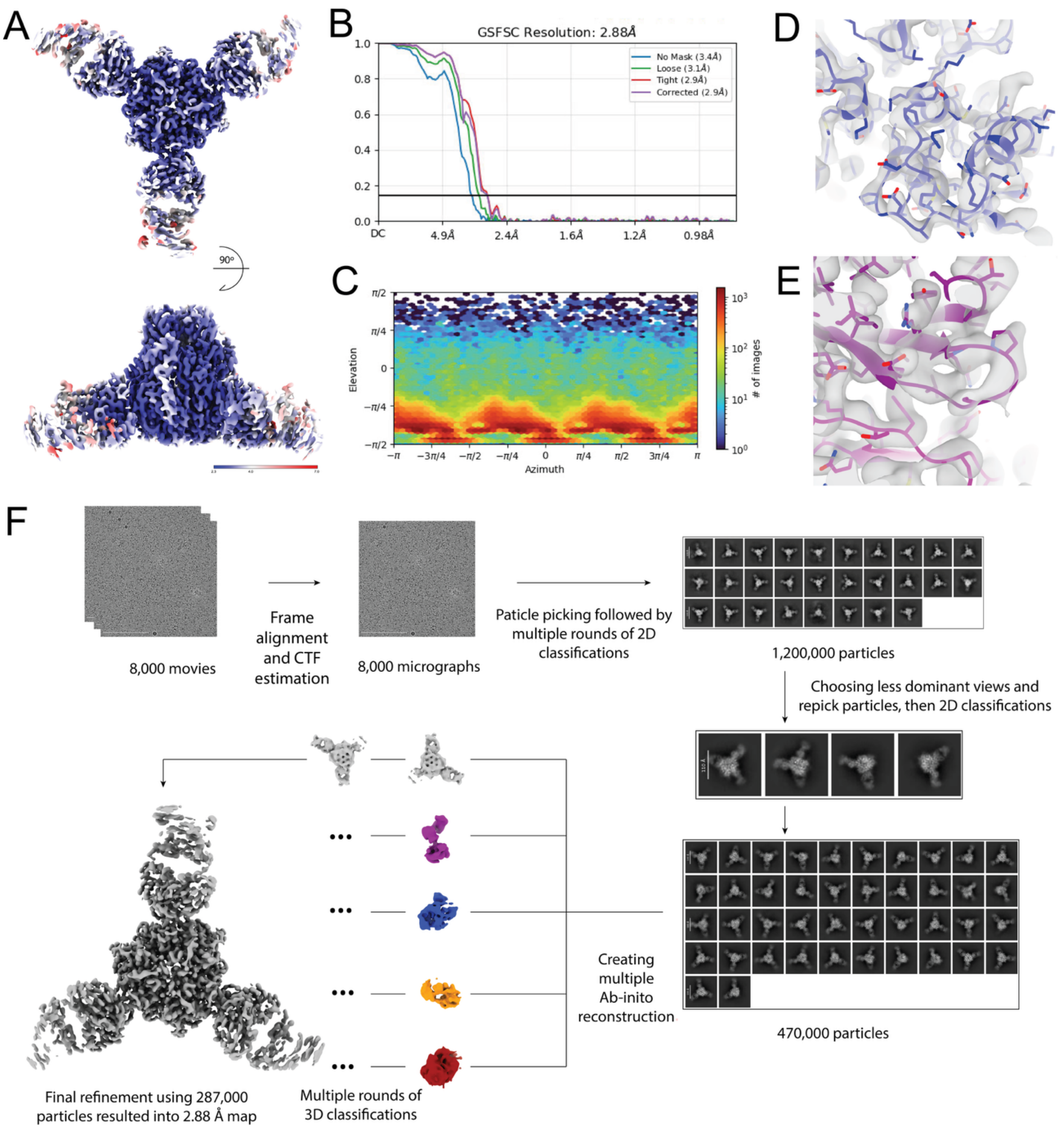


**Figure S6. MPV510 cryo-EM workflow.** (A) Local resolution map obtained for hMPV DsCav-ES2-IPDS pre-fusion F protein bound to the MPV510 Fab. (B) GSFSC curve of the refined Cryo-EM map. (C) Particle distribution map for the final refinement. (D) Model to map fit example of the hMPV DsCav-ES2-IPDS pre-fusion F protein. (E) Model to map fit example of the MPV510 Fab. (F) Overall Cryo-EM data processing workflow.
